# Integrating Green Revolution *Rht-1* Alleles in Diverse Wheat Landraces Using CRISPR/Cas9, Cytosine Base Editing and Prime Editing

**DOI:** 10.64898/2026.03.16.712109

**Authors:** Mark A Smedley, Rajani Awal, Vladimir Nekrasov, Sirisha Kaniganti, Macarena Forner, Abdul Kader Alabdullah, Simon Griffiths, Sadiye Hayta

## Abstract

The genetic diversity preserved in wheat landraces provides an important resource for broadening the genetic basis of modern wheat breeding, but its utilisation is constrained by the limited availability of effective genome-editing technologies for diverse genetic backgrounds. Here, we developed an integrated genome-engineering framework to modify *Rht-1*, a major regulator of plant height, in genetically diverse bread wheat landraces from the A.E. Watkins Collection. We combined CRISPR/Cas9-mediated deletion, cytosine base editing and prime editing to create defined *Rht-1* variants in landraces representing five underutilised ancestral groups (AG1, AG3, AG4, AG6 and AG7). CRISPR/Cas9 editing generated targeted deletions of the conserved DELLA motif while retaining the downstream GRAS domain in-frame, yielding altered *Rht-1* variants with potential utility for functional analysis and breeding. We further developed a wheat-adapted TaCBE6b-intron system and used cytosine base editing to engineer premature-stop variants in *Rht-D1*. Utilising prime editing with an adapted ePPEplus system enabled precise G-to-T conversion at the *Rht-D1* locus, generating the nucleotide substitution characteristic of the Green Revolution semi-dwarf *Rht-D1b* allele. Prime-editing activity varied substantially among Watkins genotypes, with editing detected in WATDE0394, WATDE0549, WATDE0639 and WATDE0706. At the individual-plant level, amplicon sequencing confirmed D-genome editing, with edited *Rht-D1* haplotypes reaching 100% of NGS reads in an individual WATDE0394 plant, and exceeding 50% in several plants from WATDE0394, WATDE0549 and WATDE0706. The engineered *Rht-D1* variants were associated with reduced plant height; notably, a WATDE0394 plant carrying a 100% edited D-genome haplotype reached 69 cm compared with approximately 131 cm for the control. All T1 progeny of this plant were homozygous for the prime edit, as assessed by KASP markers. Reducing plant height in these genetically diverse landraces addresses their tall architecture and associated lodging susceptibility, a practical constraint on their direct use in breeding populations. Together, these results demonstrate that complementary genome-editing technologies can generate defined *Rht-1* variation in genetically diverse wheat landraces while introducing agronomically relevant plant-height variation. The resulting founder lines provide valuable resources for functional analysis of DELLA-mediated growth regulation and a foundation for precision pre-breeding, composite cross populations, and the development of resilient wheat.

## Introduction

Wheat (*Triticum aestivum*) is one of the world’s most important staple crops, with a history of cultivation extending back in time approximately 10,000 years to the Fertile Crescent. The domestication and subsequent diversification of wheat and its ancestral species, including einkorn (*T. monococcum*) and emmer (*T. dicoccoides*), played a central role in the development of agriculture (Bilgic et al. 2016). Thousands of years of cultivation, selection and adaptation have shaped modern bread wheat, but intensive breeding has also reduced the genetic diversity represented in elite cultivars.

The A.E. Watkins Collection, assembled in the early twentieth century, comprises 827 globally sourced bread wheat landraces and represents a major reservoir of genetic diversity that remains underutilised in modern wheat breeding. Whole-genome analyses indicate that contemporary wheat cultivars capture only approximately 40% of the genetic diversity present in the Watkins Collection, with roughly 60% of the diversity represented by ancestral groups AG1, AG3, AG4, AG6 and AG7 remaining underrepresented in modern breeding programmes (Cheng et al. 2024). The Watkins landraces selected in this study represent these underutilised ancestral groups and encompass diverse geographical origins, including the Middle East (Iran and Iraq), South and Central Asia (India, Afghanistan and China), North Africa (Tunisia, Algeria and Ethiopia), southern Europe (Spain), and the Caucasus and Central Eurasia (former USSR). These accessions were selected based on their genetic diversity, geographical representation and valuable agronomic characteristics, including disease resistance, abiotic stress tolerance and other agronomically important traits. However, their tall plant architecture and associated lodging susceptibility can limit their direct deployment in modern agricultural systems.

The introduction of semi-dwarfing alleles during the Green Revolution was a major advance in wheat improvement, contributing to increased yield potential through improved lodging resistance, enhanced nitrogen responsiveness and altered assimilate partitioning (Youssefian et al. 1992). Plant height is strongly influenced by the *Reduced height-1* (*Rht-1*) homeologues, which encode DELLA proteins that are regulated by gibberellin (GA)-dependent growth. In the presence of GA, the GID1 receptor promotes interaction with DELLA proteins, triggering their ubiquitination and proteasomal degradation and thereby releasing repression of growth-promoting pathways. Mutations that reduce DELLA degradation or alter its regulatory function consequently increase growth repression and reduce stem elongation (Ross 1994).

The major Green Revolution semi-dwarfing alleles, *Rht-B1b* and *Rht-D1b*, originated from the Japanese cultivar Norin 10 and its derivatives and were subsequently incorporated into modern wheat breeding programmes (Borlaug 1968). These alleles are now present in more than 70% of current commercial wheat cultivars worldwide (Evans 1998; Han et al. 2025; Würschum et al. 2015). Both alleles contain nucleotide substitutions within the conserved LExLE region of the *Rht-1* coding sequence. The modular structure of DELLA proteins, comprising an N-terminal regulatory region with conserved DELLA and TVHYNP motifs and a C-terminal GRAS domain, provides unique opportunities to investigate how individual protein regions regulate growth (Fig. 1A). In the semi-dwarfing alleles *Rht-B1b* and *Rht-D1b*, specific single-nucleotide substitutions disrupt this structure. In *Rht-B1b*, a C-to-T substitution converts the glutamine codon CAG to the amber stop codon TAG, whereas in *Rht-D1b*, a G-to-T substitution converts the glutamic acid codon GAG to TAG (Peng et al., 1999; Van De Velde et al., 2021). Although these mutations involve different nucleotide conversions, both alleles generate premature stop codons that can promote translational reinitiation downstream of the mutation site (Fig. 1C). This process produces N-terminally truncated DELLA proteins that retain the functional C-terminal GRAS domain but lack the essential N-terminal GA-responsive regulatory region. The resulting alteration in DELLA protein structure and function leads to enhanced, constitutive repression of GA-dependent growth, ultimately contributing to the characteristic GA-insensitive semi-dwarf phenotype (Chen & Liu, 2023; Kowalski et al., 2016; Pearce et al., 2011; Thomas, 2017; Van De Velde et al., 2021).

**Figure 1.**
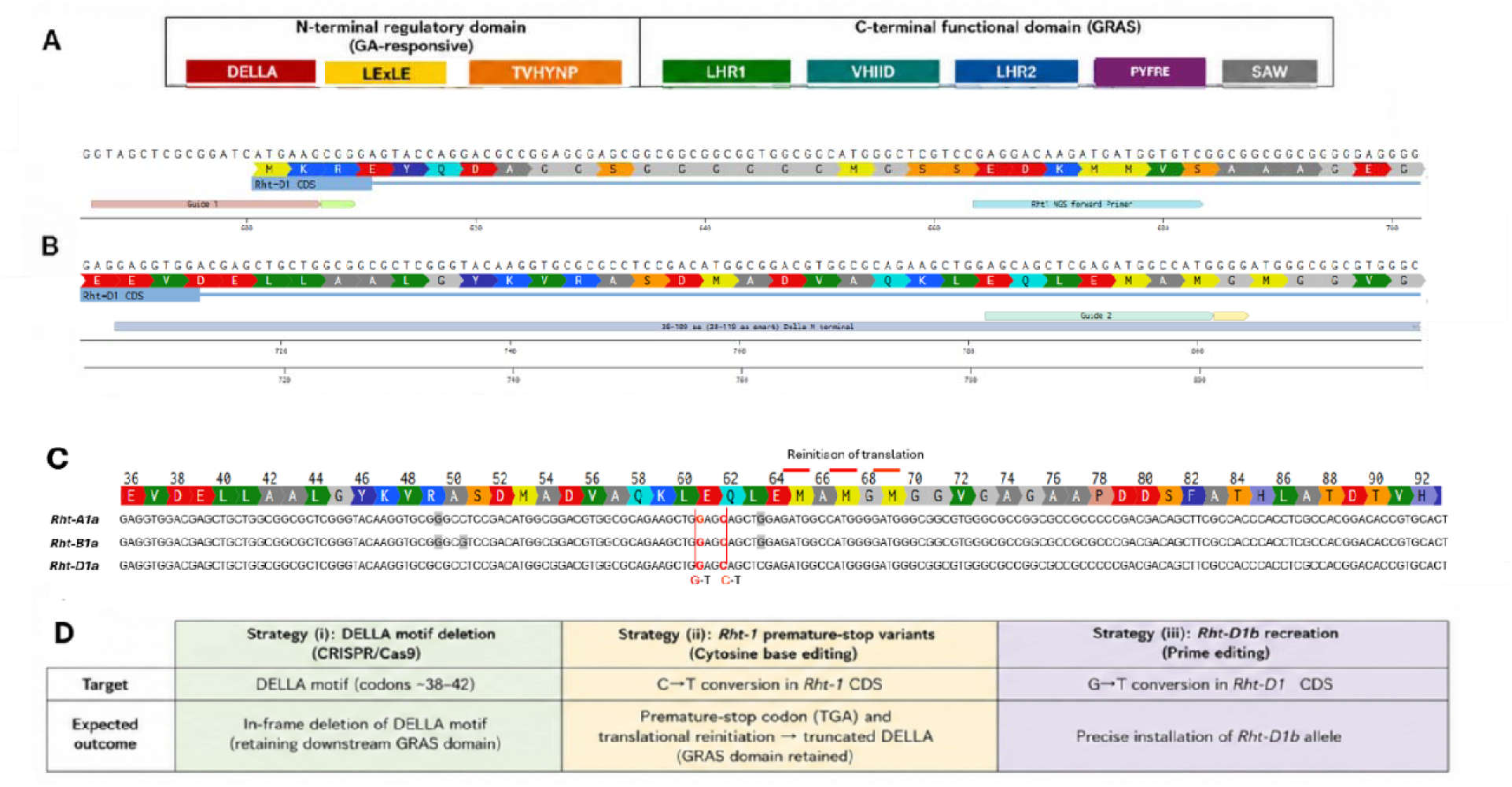
Domain architecture and genome-editing strategies targeting wheat *Rht-1* homeologs. **(A)** Domain architecture of the wheat *Rht-1* homeologs. Conserved N-terminal motifs, including DELLA, LExLE and TVHYNP, and the C-terminal GRAS domain, comprising the LHR1, VHIID, LHR2, PYFRE and SAW motifs, are indicated. **(B)** *Rht-D1a* sequence showing the positions of the gRNAs used to delete the conserved DELLA motif while maintaining the downstream GRAS domain in frame. Red lines indicate the expected Cas9 cleavage sites. **(C)** Alignment of nucleotide sequences corresponding to the N-terminal coding regions of the *Rht-1* homeologs. Target nucleotide substitutions are highlighted, including the C→T conversion used to introduce a premature stop codon and the G→T conversion used to generate the *Rht-D1b* allele through introduction of the corresponding premature stop codon. Red underlines indicate the predicted reinitiation sites. **(D)** Schematic summary of the genome-editing strategies used in this study, including CRISPR/Cas9-mediated deletion of the conserved DELLA motif, cytosine base editing to introduce a premature stop codon, and prime editing to generate the *Rht-D1b* allele.

The defined nucleotide changes underlying *Rht-B1b* and *Rht-D1b* provide particularly suitable targets for precision genome engineering. The C-to-T substitution required to recreate *Rht-B1b* is compatible with cytosine base editing, whereas the G-to-T substitution required for *Rht-D1b* cannot be generated directly using conventional cytosine or adenine base-editing systems and therefore requires an alternative editing strategy such as prime editing. Combining complementary genome-editing technologies consequently provides a route to recreate established agronomically important alleles while also generating new *Rht-1* variants in diverse genetic backgrounds.

Recent advances in genome engineering have created new opportunities to recreate naturally occurring alleles and generate defined functional variation directly within elite and non-elite germplasm. Unlike conventional breeding, which relies on existing allelic variation and repeated crossing, precision genome editing enables targeted modification of individual nucleotides or defined protein domains while preserving the broader genetic background of valuable landraces. CRISPR/Cas9 can generate targeted mutations and deletions, whereas cytosine base editors enable programmable C•G-to-T•A conversion without requiring double-strand DNA breaks. Prime editing further expands the range of possible sequence changes by enabling precise installation of nucleotide substitutions and small sequence alterations through a reverse-transcription-based mechanism (Anzalone et al. 2019).

In this study, we adapted complementary editing technologies for application in diverse wheat germplasm. We adapted CBE6b (Zhang et al. 2024) through codon optimisation and intron incorporation to establish a TaCBE6b-intron system capable of generating targeted C-to-T substitutions in wheat. We also adapted the enhanced plant prime editor plus (ePPEplus), comprising an nCas9–M-MLV reverse transcriptase fusion and incorporating the Csy4 RNA-processing system, for use within our established wheat transformation platform (Ni et al., 2023). Advances in editor architecture, pegRNA design, reverse-transcriptase optimisation and modulation of DNA repair pathways have further improved the potential of these technologies for precise genome engineering in plants and complex crop genomes (Chen and Liu 2023; Liu et al. 2021).

Here, we develop an integrated genome-engineering framework to generate and characterise *Rht-1* allelic variation in wheat landraces representing the previously underutilised Watkins ancestral groups AG1, AG3, AG4, AG6 and AG7. We employed three complementary editing strategies: (i) CRISPR/Cas9-mediated deletion of the conserved DELLA motif while retaining the downstream GRAS domain in-frame (Fig. 1B); (ii) cytosine base editing to introduce defined C-to-T substitutions and generate premature-stop *Rht-1* variants, including premature-stop variants in *Rht-D1*; and (iii) prime editing to precisely install the G-to-T substitution characteristic of the semi-dwarf *Rht-D1b* allele (Fig. 1D).

Together, these approaches establish an expanded precision genome-editing toolbox for genetically diverse wheat germplasm and generate molecularly defined *Rht-1* variants in underutilised landrace backgrounds, including precise engineering of the *Rht-D1b* allele. The resulting founder lines combine engineered growth-regulatory variation with diverse ancestral genetic backgrounds and, where associated with reduced plant height, address the tall architecture and lodging susceptibility that can limit the direct use of these landraces in breeding populations. These materials provide resources for functional dissection of DELLA-mediated GA responses and a foundation for precision pre-breeding, composite cross populations and the development of climate-resilient wheat.

## Material and Methods

### Plant Material and Growth Conditions

The spring wheat (*Triticum aestivum*) cultivar Paragon and 42 Watkins wheat landraces representing the genetic diversity of the A.E. Watkins Collection were used in this study. The Watkins accessions comprised WATDE0005, WATDE0008, WATDE0031, WATDE0040, WATDE0042, WATDE0047, WATDE0051, WATDE0052, WATDE0057, WATDE0124, WATDE0150, WATDE0156, WATDE0158, WATDE0185, WATDE0215, WATDE0216, WATDE0383, WATDE0385, WATDE0394, WATDE0401, WATDE0415, WATDE0442, WATDE0446, WATDE0451, WATDE0452, WATDE0481, WATDE0549, WATDE0553, WATDE0555, WATDE0574, WATDE0585, WATDE0588, WATDE0592, WATDE0611, WATDE0616, WATDE0626, WATDE0639, WATDE0643, WATDE0653, WATDE0695, WATDE0706 and WATDE0895.

Gene-editing experiments included CRISPR/Cas9-mediated targeted mutagenesis, prime editing and cytosine base editing. CRISPR/Cas9-mediated targeted mutagenesis was initially established in 20 selected Watkins landraces to assess genome-editing efficiency across diverse genetic backgrounds. Prime editing was subsequently optimised and expanded across all 42 Watkins landraces. Cytosine base-editing experiments were performed using the spring wheat cultivar Paragon.

Individual seeds were sown in 11-cm-diameter pots containing a cereal compost mixture comprising peat, sterilised loam and horticultural grit, supplemented with limestone, base fertiliser, slow-release fertiliser and a wetting agent. Plants were vernalised at 5 °C under an 8-h photoperiod for 4 weeks for spring accessions or 8 weeks for winter accessions, with a light intensity of 70 μmol m⁻² s⁻¹.

Following vernalisation, plants were transferred to controlled-environment rooms (CERs) maintained at 18 ± 1 °C during the day and 15 ± 1 °C at night, with 65% relative humidity and a light intensity of approximately 350 μmol m⁻² s⁻¹. Plants were maintained under automated irrigation as previously described by Hayta et al. (2021).

After a further two weeks of growth, seedlings were transplanted into 2-L pots containing the same cereal compost mixture described above. To ensure a continuous supply of immature embryos for transformation experiments, seeds from different Watkins accessions were sown in small batches at weekly intervals. Paragon, a spring wheat cultivar that does not require vernalisation, was grown directly in controlled-environment rooms without a vernalisation treatment.

### *Rht-D1 d*ecoupling N terminal DELLA from C terminal GRAS construct preparation

Level 2 CRISPR/Cas9 plasmids were developed based on the pGoldenGreenGate (pGGG) system (Smedley et al., 2021). These vectors contain a hygromycin resistance marker, a heat-shock–inducible Cre–LoxP–excisable GRF4–GIF1 regeneration enhancer, and an intron-enhanced Cas9 expression cassette, and allow the insertion of guide RNAs (gRNAs) driven by the wheat TaU6 promoter.

Briefly, a level 1 expression cassette comprising the rice ubiquitin promoter (OsUbi) driving a *Zea mays*–codon-optimised Cas9 containing 13 introns (Cas9-int) (Grützner et al., 2021), flanked by 5T and 3T nuclear localisation signals (NLS) and terminated by the NOS terminator, was assembled using standardised Golden Gate cloning with *Eco*31L.

For level 2 assembly, a modified binary pGGG vector lacking the *lacZ* bacterial selection cassette, termed pGGG (minus), was used. This vector accepts level 1 modules via BpiI digestion. The Pv*Ubi2*::*hpt* selectable marker, the heat-shock–inducible Cre–LoxP Zm*Ubi*::*GRF4–GIF1* cassette, the level 1 *lacZ* acceptor module and the Os*Ubi*::Cas9-int cassette were assembled into pGGG (minus) using BpiI. Correct assemblies were identified by blue colony selection and whole plasmid Next Generation Sequencing (NGS).

To facilitate visual screening, a chromoprotein bacterial marker cassette (amilCP, purple) was designed and inserted into the acceptor region of the pGGG backbone using *Eco*31L. This configuration allows the subsequent insertion of level 1 components, including gRNA arrays, at MoClo golden gate positions 1–7. The resulting vector was designated pGGG (S1) Purple.

Two gRNAs targeting the *Rht1* locus were designed to specifically target the D-genome homeolog, based on the wheat reference gene models *TraesCS4A02G271000*, *TraesCS4B02G043100*, and *TraesCS4D02G040400* (Table 1). The final pGGG (S1) construct contained two TaU6-driven gRNAs targeting *Rht1*, the hygromycin selectable marker, the heat-shock–excisable GRF4–GIF1 cassette, and the intron-enhanced Cas9 driven by the rice ubiquitin promoter. After full plasmid sequencing, the completed pGGG-Rht1 vector (Fig 2A) was electroporated into *Agrobacterium* stain AGL1 and standard inoculums made as described in (Hayta et al., 2019).

**Figure 2.**
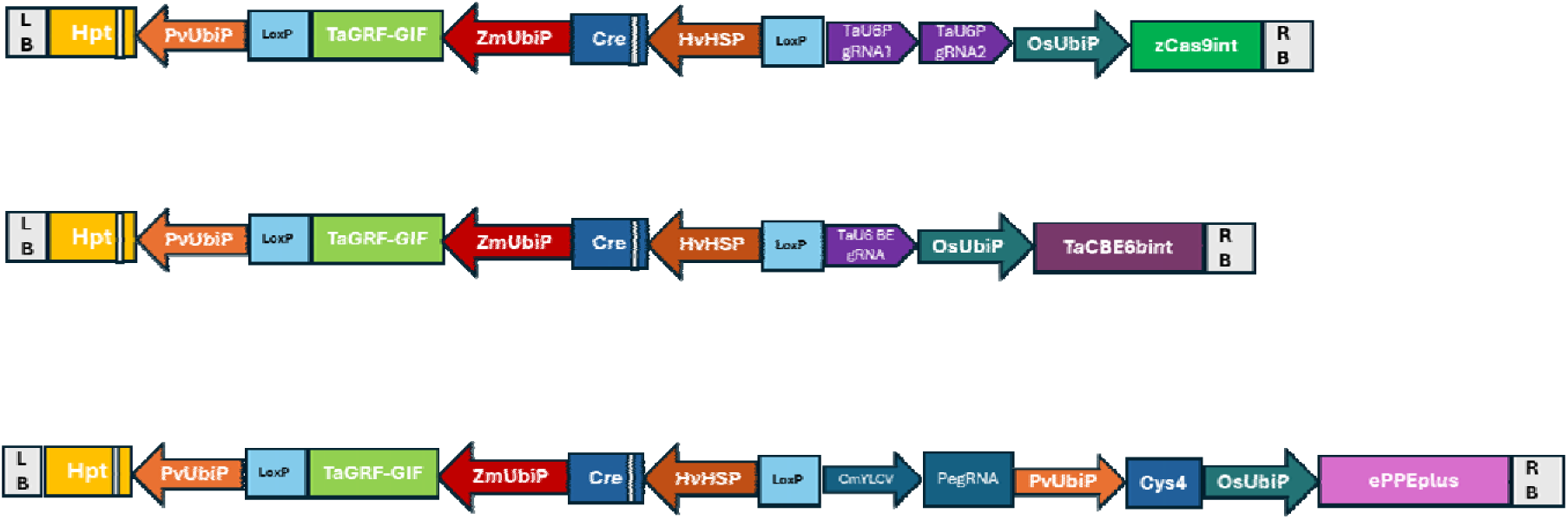
Schematic representation of genome editing constructs used in this study. (**A**) The pGoldenGreenGate (S1) CRISPR/Cas9 construct targeting Rht1, containing two guide RNAs driven by the TaU6 promoter and an intron-enhanced Cas9 nuclease driven by the rice ubiquitin (OsUbi) promoter. (**B**) The TaCBEint-Rht1 base editing construct, containing an intron-enhanced, wheat codon-optimised cytosine base editor TaCBE6b driven by the rice ubiquitin promoter, together with a TaU6-driven guide RNA targeting Rht-D1. (**C**) The pGGG-ePPEplus-Rht1 prime editing construct, containing an Rht1-targeting pegRNA cassette with the reverse transcriptase template sequence, the enhanced plant prime editor (ePPE) comprising a Cas9 nickase (nCas9) fused to an engineered Moloney murine leukaemia virus (M-MLV) reverse transcriptase, and the Csy4-mediated guide RNA processing system. (All constructs contained a hygromycin resistance selectable marker and a heat-shock-inducible Cre– LoxP excisable GRF4–GIF1 module to enhance transformation efficiency and enable subsequent removal of the regeneration-associated cassette).

**Table 1.**
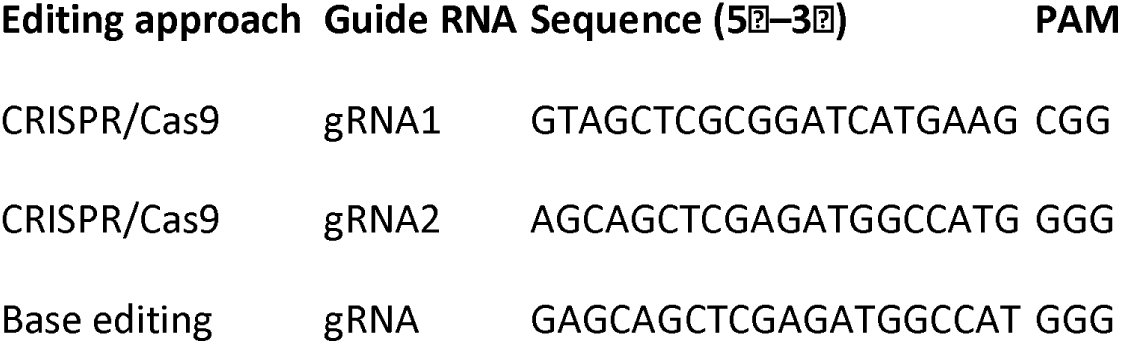
Guide RNAs used for CRISPR, and base editing of the *Rht1* locus.

### Base Editing Construct

To improve gene expression and increase the translational efficiency of CBE6b the sequence of CBE6b was first codon optimised for expression in wheat and intron enhanced by including 7 introns within the coding sequence. Wheat codon optimisation of CBE6b was carried out using the online tool at (www.Genewiz.com) and denoted as TaCBE6b. The splice site prediction online tool Spliceator (www.lbgi.fr/spliceator/) was used to predict potential splice sites (Scalzitti et al. 2021), and canonical sites were chosen to insert intronic sequences as described in Grützner et al. (2021). The TaCBE6b-intron was synthesised as a golden gate MoClo level 0 component, internal *Eco*31L and BpiI restriction enzyme sites were removed by silent mutation and standardised MoClo overhangs and *Eco*31L sites added for assembly. Construct assembly was performed using MoClo golden gate assembly (Weber et al., 2011), into a Level 2 vector based on the binary vector pGoldenGreenGate (pGGG) (Smedley et al., 2021). The pGGG (V) vector contains the hygromycin selectable marker (hpt) and Cat1 intron driven by the switchgrass ubiquitin promoter, a Cre recombinase heat shock excisable GRF4-GIF1 gene fusion driven by the maize ubiquitin promoter and an acceptor cassette which allows insertion of level 1 golden gate components via *Bpi*I. The TaCBE6b-intron synthon was assembled using *Eco*31L as a Level 1 transcriptional cassette under the control of the rice ubiquitin promoter and the nos terminator. The wheat Level 1 TaU6 guide RNA acceptor (Addgene # 165599) and the Level 1 TaCBE6b-intron transitional cassette were both assembled into the Level 2 pGGG (V) vector using *Bpi*I. The resulting vector was deemed pGGG TaCBEint single guide acceptor (Fig 2B).

CBE6b exhibits an activity window within the protospacer spanning positions 3 to 13 from the 5T end, with optimal cytosine deamination occurring between positions 4 and 8. A base editing guide RNA (BE-gRNA) (Table 1) was therefore designed to position the targeted cytosine within this optimal editing window to maximise editing efficiency and to recreate the early stop codon (Fig 3A). The BE-gRNA was ordered as two complimentary oligos and assembled into pGGG TaCBEint single guide acceptor via *Eco*31L.

**Figure 3.**
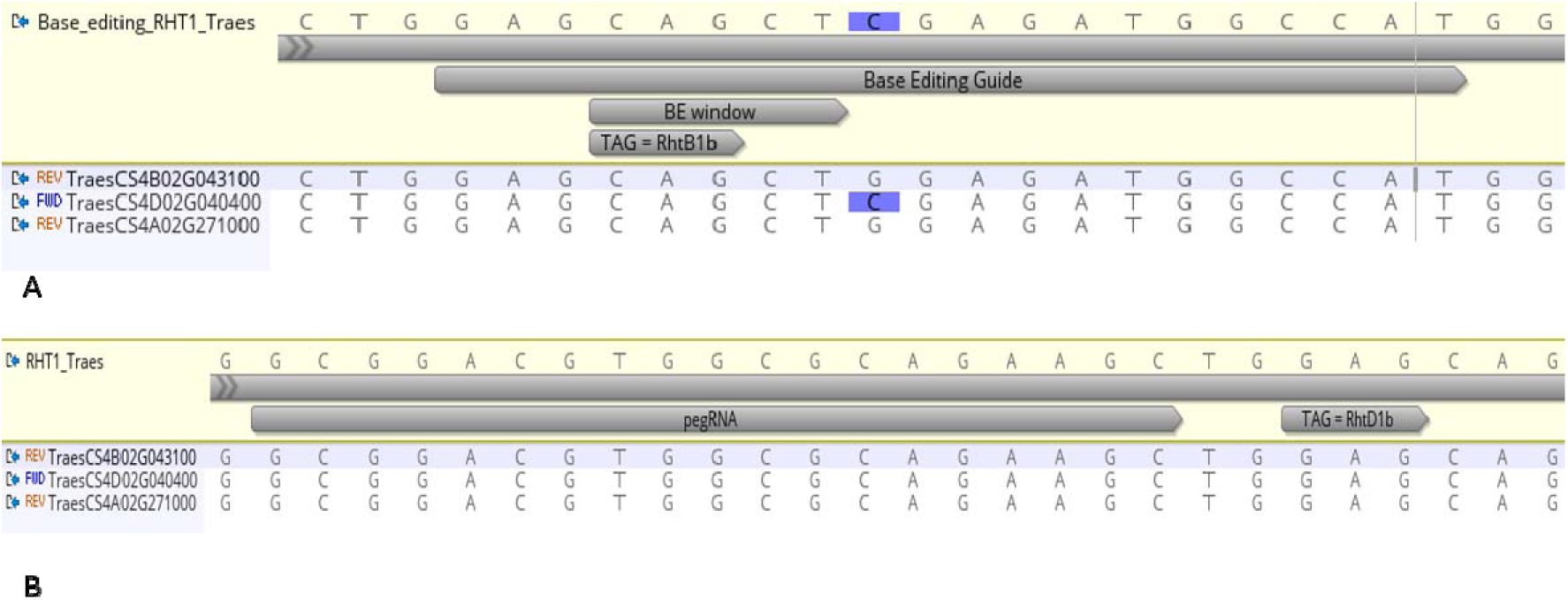
Design of base editing and prime editing strategies for generating semi-dwarfing Rht-1 alleles. (**A**) Alignment of the three wheat Rht-1 homeologs showing the base editing guide RNA (gRNA) designed to recreate the Rht-1 early stop codon mutation in D subgenome. (**B**) Schematic representation of the wheat genomic target region and prime editing guide RNA (pegRNA) design used to introduce the Rht-D1b early stop codon mutation.

### Prime Editing Construct

The pGGG ePPEplus–Rht1 vector was constructed to enable prime editing of the *Rht1* locus. This construct contains a prime editing guide RNA (pegRNA) encoding both the spacer sequence and the reverse transcription (RT) template, together with the enhanced plant prime editor plus (ePPEplus), consisting of an nCas9–M-MLV reverse transcriptase fusion (Ni et al., 2023). The vector also incorporates the Csy4 RNA processing system to enable precise maturation of pegRNAs from polycistronic transcripts.

The prime editing target was identified by eye (Fig. 3B) and the PlantPegDesigner online tool http://www.plantgenomeediting.net (Jin et al. 2023) used to design the pegRNA with the desired template for the reverse transcriptase (Supplementary Fig 1).

Construct assembly was performed using MoClo golden gate assembly (Weber et al., 2011). For level 1 assembly, the pegRNA was synthesised flanked by Csy4 recognition sites and 4 bp overhangs plus *Eco*31L recognition sites to enable insertion downstream of the Cestrum yellow leaf curling virus (CmYLCV) promoter to drive expression. The ePPEplus CDS was assembled as a level 1 expression cassette driven by the rice ubiquitin promoter and nos terminator, and the Cys4 level 1 expression cassette assemble with the switchgrass ubiquitin promoter (PvUbiP) driving expression, both via *Eco*31L. The Level 1 expression cassettes were assembled into our preferred Level 2 binary vector pGoldenGreenGate-M (pGGG-M) Addgene #165422 via *Bpi*I (Smedley et al. 2021) along with the hygromycin selectable marker (*hpt*) and the GRF4-GIF1 fusion Addgene #198046. The completed vector pGGG ePPEplus Rht1 (Fig. 2c) was whole plasmid sequenced to confirm authenticity before electroporation into the hypervirulent *Agrobacterium* tumefaciens strain AGL1 (Lazo et al. 1991) and standard inoculums prepared as described in Hayta et al. (2019) for wheat transformation.

### *Agrobacterium*-mediated Transformation across Watkins Lines

*Agrobacterium*-mediated transformation was performed as previously described by Hayta et al. (2021, 2026), with minor modifications to optimise transformation and regeneration of the Watkins landraces. Following co-cultivation, transformed calli were selected on media containing hygromycin B. Selection was carried out using 5 mg L⁻¹ hygromycin B during the first selection stage (S1) and 10 mg L⁻¹ during the second selection stage (S2).

Selected calli were transferred to regeneration medium (R1) containing 10–15 mg L⁻¹ hygromycin B and 2.5 mg L⁻¹ zeatin riboside. Cultures were subsequently subcultured onto fresh regeneration medium every two weeks through three regeneration stages (R1–R3) to promote shoot development while maintaining selection pressure. Regenerated shoots were transferred to rooting medium containing 20 mg L⁻¹ hygromycin B.

These selection and regeneration conditions were applied consistently throughout the transformation procedure, with minor adjustments made where necessary to optimise regeneration of individual Watkins landraces.

### Copy Number Analysis

Leaf samples (0.5 to 0.7 cm) were collected, and DNA was extracted using a protocol adapted from (Pallotta et al., 2003) by JIC Genotyping Facility. Transgenesis was confirmed, and transgene copy number determined using TaqMan qPCR with probes as described by Hayta et al. (2019). Copy number values were calculated according to the comparative Delta Ct (Δ Ct) method (Livak & Schmittgen, 2001).

### PCR screening and Targeted Amplicon Sequencing

PCR reactions were performed using ReadyMix™ Taq PCR Reaction Mix (Sigma, cat. no. P4600-100RXN) in a total volume of 20 µL. Each reaction contained 10 µL PCR ReadyMix™, 2 µL (∼100 ng) of wheat genomic DNA, 1 µL of each forward and reverse primer (10 µM), and sterile laboratory-grade water to a final volume of 20 µL.

For transgene screening, a 335 bp fragment of the hygromycin phosphotransferase (*hpt*) gene was amplified using primers HygF (5T-AGGCTCTCGATGAGCTGATGCTTT-3T) and HygR (5T-AGCTGCATCATCGAAATTGCCGTC-3T). PCR amplification was performed with an initial denaturation at 95 °C for 3 min, followed by 34 cycles of 95 °C for 30 s, 58 °C for 30 s, and 72°C for 1 min, with a final extension at 72 °C for 7 min and a final hold at 10 °C. PCR products were separated on a 1% agarose gel containing ethidium bromide (1 µg mL⁻¹) and visualised under UV illumination.

For targeted amplicon sequencing, the *Rht1* DELLA target region was amplified using gene-specific primers containing illumina NGS adapter sequencing tails (Table 2). PCR amplification was performed using an initial denaturation at 95 °C for 3 min, followed by 35 cycles of 95 °C for 30 s, 58 °C for 30 s, and 72 °C for 25 s, followed by a final extension at 72 °C for 7 min and a final hold at 10 °C. The expected amplicon sizes were 455 bp for the unedited *Rht1* DELLA allele and 326 bp for *Rht1* PE/BE alleles. The PCR products were pooled and sequenced using an Illumina next-generation sequencing platform at the QIB DNA Sequencing Facility. Sequencing data were analysed to identify and quantify editing outcomes.

**Table 2.**
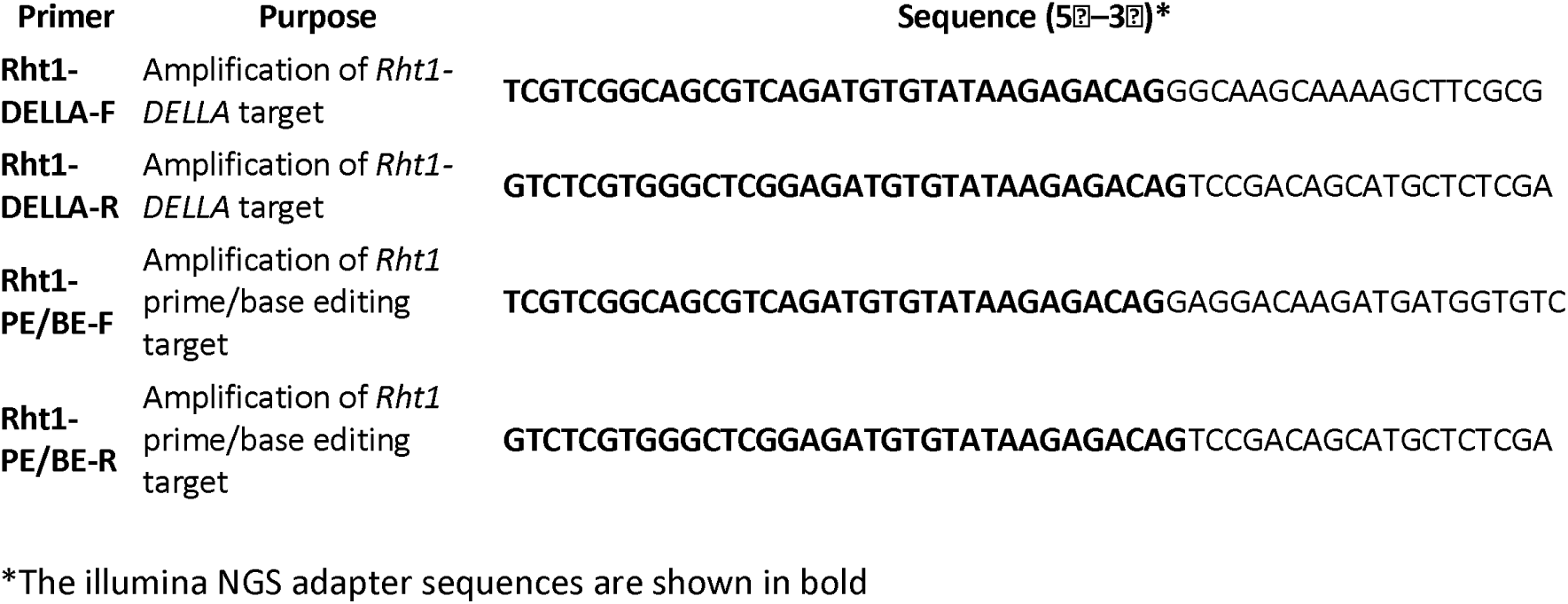
Illumina-tailed primers used for amplicon sequencing of *Rht1* editing events.

### Targeted Amplicon Sequencing Data Analysis

Sequencing reads were mapped to the appropriate reference sequence using BWA v0.7.17. The resulting alignment files were converted to BAM format, sorted, and indexed using SAMtools v1.10. Processed BAM files were analysed using Stack Mapping Anchor Points (SMAP) to identify unique haplotypes (Develtere et al. 2023), estimate their relative frequencies, and characterise CRISPR/Cas9-induced sequence variation, including insertions, deletions, and nucleotide substitutions.

Plants were classified as edited when CRISPR/Cas9-derived haplotypes accounted for ≥30% of sequencing reads at the target locus. Haplotype frequency distributions were used to classify editing outcomes as homozygous, heterozygous, or mosaic based on the relative abundance of edited and wild-type alleles. Sequence alignments and selected editing events, including large deletions, were manually inspected using Integrative Genomics Viewer (IGV).

Editing efficiency was calculated as the percentage of T₀ plants containing at least one confirmed CRISPR/Cas9-induced mutation relative to the total number of T₀ plants analysed for each independent transformation experiment.

Large deletion alleles identified from SMAP analysis were further selected and visualised using Benchling to confirm the predicted genomic modifications. Representative in-frame deletion alleles were characterised based on removal of the DELLA and TVHYNP motifs while maintaining the open reading frame and preserving the C-terminal GRAS domain, consistent with the predicted generation of GA-insensitive DELLA variants.

Following the identification of Prime edited T₀ plants, the resulting T₁ progeny were further genotyped to identify and confirm the inheritance of the engineered *Rht-D1b* allele. KASP genotyping was performed by the John Innes Centre Genotyping Service using the co-dominant wMAS00002 SNP marker. The assay distinguishes the wild-type G allele (FAM) from the dwarfing T allele (VIC). Genotypes were assigned using SNPviewer as homozygous wild type (G/G), heterozygous (G/T), or homozygous dwarf (T/T).

### Plant Height Response to Applied GA_3_

To assess gibberellin responsiveness of Rht-1 prime edited wheat lines, homozygous, heterozygous and WT T₁ progeny were grown under controlled environmental conditions until the three-leaf stage. Gibberellic acid (10 µM) (GA₃; Sigma-Aldrich, G7645) was dissolved in ethanol and diluted with distilled water to prepare the working solution. Plants were sprayed with 5 mL of GA₃ solution per seedling until the leaf surfaces were uniformly wetted. Control plants received a mock treatment. Treatments were applied every 3 days, in total plants received 6 sprays. Plant growth responses were monitored throughout the treatment period.

Plant height was recorded before and after GA₃ treatment, measured from the soil surface to the leaf tip. The relative GA response was calculated by comparing the change in plant height following GA₃ treatment between edited lines and untreated/mock-treated controls. Eight biological replicates were analysed for each genotype.

Statistical analysis was performed using a Type III factorial ANOVA to assess the effects of genotype, GA₃ treatment, and their interaction on plant height response. Where significant effects were detected, appropriate post hoc multiple-comparison tests were performed. Statistical significance was defined as *P* < 0.05.

## Results

### Transformation of Watkins Wheat Landraces Using the TaGRF4–GIF1 Regeneration System

This study represents the first successful *Agrobacterium*-mediated transformation of Watkins wheat landraces, providing access to a genetically diverse resource that has remained largely underutilised for functional genomics because many landraces are recalcitrant to transformation. Using an *Agrobacterium*-mediated transformation system incorporating the Ta*GRF4–GIF1* developmental regulator fusion (Debernardi *et al*., 2020), transgenic plants were successfully recovered from 13 Watkins landraces representing four ancestral groups (AG1, AG3, AG6 and AG7).

Substantial variation was observed in transformation performance among the Watkins accessions (Table 3). A total of 90 independent transgenic plants were recovered, with the highest numbers obtained from WATDE0585 (26 plants), WATDE0385 (18 plants), WATDE0005 (14 plants), and WATDE0401 (13 plants). In contrast, only one or two transgenic plants were recovered from several accessions, including WATDE0150, WATDE0574, WATDE0040, WATDE0643, WATDE0008 and WATDE0442, whereas no transgenic plants were recovered from WATDE0185. These findings suggest that the *TaGRF4–GIF1* system can facilitate transformation across a broader range of wheat genotypes, although differences in transformation and regeneration responses remain apparent among genotypes, consistent with the inherent genotype dependence of wheat transformation.

**Table 3.** CRISPR/Cas9 editing efficiencies in Watkins wheat landraces in *Rht1* Della deletion.

| **Watkins line** | **Ancestral group** | **Transgenic plants recovered** | **Plants screened** | **Edited plants** | **Editing efficiency (%)** |
| --- | --- | --- | --- | --- | --- |
| WATDE0005 | AG3 | 39 | 39 | 33 | 85 |
| WATDE0156 | AG6 | 3 | 3 | 3 | 100 |
| WATDE0385 | AG3 | 18 | 15 | 10 | 56 |
| WATDE0451 | AG7 | 7 | 16 | 15 | 75 |
| WATDE0585 | AG1 | 26 | 30 | 21 | 70 |
| WATDE0401 | AG1 | 13 | 26 | 17 | 65 |
| WATDE0150 | AG6 | 1 | 1 | 1 | 100 |
| WATDE0185 | AG6 | 1 | 1 | 0 | 0 |
| WATDE0574 | AG1 | 1 | 9 | 9 | 100 |
| WATDE0643 | AG4 | 2 | 2 | 2 | 100 |
| WATDE0442 | AG6 | 2 | 2 | 1 | 50 |
| WATDE0008 | AG7 | 4 | 4 | 4 | 100 |
| WATDE0446 | AG6 | 2 | 2 | 2 | 100 |

The successful recovery of transgenic plants across diverse Watkins landraces highlights the potential of the *TaGRF4–GIF1* regeneration system to overcome genotype-associated barriers to wheat transformation. Debernardi et al. (2020) demonstrated that expression of the *TaGRF4–GIF1* chimera substantially enhances regeneration capacity in wheat, achieving transformation efficiencies of up to 77.5% in the highly transformable spring wheat cultivar Fielder and extending transformation to previously recalcitrant cultivars. Developmental regulators such as *GRF4–GIF1* are thought to enhance cellular competence for regeneration by promoting the proliferation and developmental plasticity of immature embryogenic cells. Importantly, incorporation of this technology into the transformation pipeline can reduce genotype-dependent limitations and broaden the developmental window during which competent explant material can successfully regenerate (Hayta et al. 2026). This is particularly valuable in wheat, where transformation efficiency is strongly influenced by genotype, embryo developmental stage, and tightly controlled growth conditions. Thus, the use of *TaGRF4–GIF1* provides a robust strategy for accelerating transformation and enabling efficient recovery of transgenic plants from genetically diverse wheat germplasm.

Despite the efficiency gains offered by *GRF4–GIF1*, constitutive expression can cause pleiotropic effects, including reduced fertility, decreased spikelet number, and alterations in plant architecture, with the severity of these effects varying among genotypes. To minimise potential developmental effects associated with prolonged expression of the regeneration-promoting regulator, we applied a heat-inducible Cre/lox excision system to remove the *GRF4–GIF1* cassette following plant regeneration, as previously reported by Hayta *et al*. (2026). We confirmed successful excision of the *GRF4–GIF1* cassette after plant regeneration, thereby restricting its activity to the regeneration phase and limiting its effects during subsequent plant development. This post-regeneration excision strategy therefore provides an effective means of reducing the potential pleiotropic effects associated with prolonged *GRF4–GIF1* expression, including effects on fertility and seed production.

Despite the improved regeneration capacity conferred by Ta*GRF4–GIF1*, considerable variation in transformation performance remained among the Watkins landraces. Such genotype-dependent responses are well documented in wheat and are likely to reflect the extensive genetic diversity represented within the Watkins Collection. Although donor plants were grown for a relatively short and controlled period to minimise environmental variation, differences in embryo quality, developmental stage, and intrinsic regenerative capacity among genotypes are likely to have contributed to the observed differences. In addition, the Watkins landraces originate from geographically diverse environments and have undergone adaptation to distinct climatic and agronomic conditions. Consequently, differences in growth rate, developmental timing, and physiological status during donor plant production may influence immature embryo quality and, subsequently, transformation and regeneration outcomes. Similar genotype-dependent effects have been reported across both elite wheat cultivars and diverse wheat germplasm collections (Hayta et al. 2026).

Importantly, the recovery of transgenic plants from multiple Watkins landraces demonstrates that *TaGRF4–GIF1*-assisted transformation can provide a practical route to accessing genetically diverse wheat germplasm that has historically been recalcitrant to transformation and, has not previously been transformed. This expands the experimental scope of functional genomics beyond a relatively small number of transformation-compatible wheat cultivars and provides a foundation for applying genome editing and other precision-breeding approaches directly in diverse Watkins genetic backgrounds. The establishment of transformation competence across these landraces therefore represents an important step towards exploiting the substantial, previously underutilised genetic diversity of the Watkins Collection for gene discovery, functional validation, and precision breeding.

### CRISPR Editing Efficiencies Across Watkins Wheat Landraces

Gene editing efficiencies were evaluated by targeted amplicon sequencing of primary transgenic (T₀) plants generated following *Agrobacterium*-mediated transformation of Watkins wheat landraces. Across all experiments, a diverse collection of CRISPR/Cas9-induced alleles were identified, demonstrating efficient editing of the target locus and a broad spectrum of DNA repair outcomes.

Sequence analysis revealed numerous distinct haplotypes containing small insertions and deletions (indels), as well as larger deletions centred on the predicted Cas9 cleavage site. Both frameshift and in-frame mutations were recovered. Frameshift mutations were predicted to introduce premature stop codons, resulting in truncated DELLA proteins. In contrast, several in-frame alleles precisely removed the conserved DELLA and TVHYNP motifs while maintaining the reading frame and preserving the C-terminal GRAS domain. The DELLA and TVHYNP motifs are essential for recognition by the GA-bound GID1 receptor complex, which triggers DELLA degradation and promotes GA-responsive growth. Their removal is therefore predicted to abolish GID1-mediated DELLA degradation while retaining the transcriptional regulatory activity conferred by the C-terminal GRAS domain, resulting in constitutively stable protein, a GA-insensitive phenotype, and reduced plant height (dwarfism).

Editing efficiencies varied among Watkins landraces but were generally high (Table 3). This finding is consistent with the observations of Grützner et al. (2021) who demonstrated that the incorporation of 13 introns into a plant- or maize-codon-optimised Cas9 coding sequence, generating an “intronised” Cas9 (zCas9i), substantially increased CRISPR/Cas9 genome-editing efficiency in *Arabidopsis thaliana*. A total of 150 T₀ plants representing 13 Watkins landraces were analysed, of which 118 plants carried CRISPR/Cas9-induced mutations, corresponding to an overall editing efficiency of 78.7%. Editing efficiencies ranged from 0% to 100%. Complete editing (100%) was achieved in WATDE0156, WATDE0150, WATDE0574, WATDE0643, WATDE0008 and WATDE0446, while WATDE0451 and WATDE0585 exhibited editing efficiencies of 75% and 70%, respectively. The largest population analysed, WATDE0005 (71 T₀ plants), showed a high editing efficiency of 85% (Fig 4A), whereas WATDE0385 displayed a lower editing efficiency of 56%. No edited plants were recovered from WATDE0185, for which only a single T₀ transgenic event was obtained. Edited plants were recovered from landraces representing the ancestral groups AG1, AG3, AG4, AG6 and AG7, demonstrating that efficient CRISPR/Cas9-mediated mutagenesis can be achieved across genetically diverse wheat backgrounds

**Figure 4.**
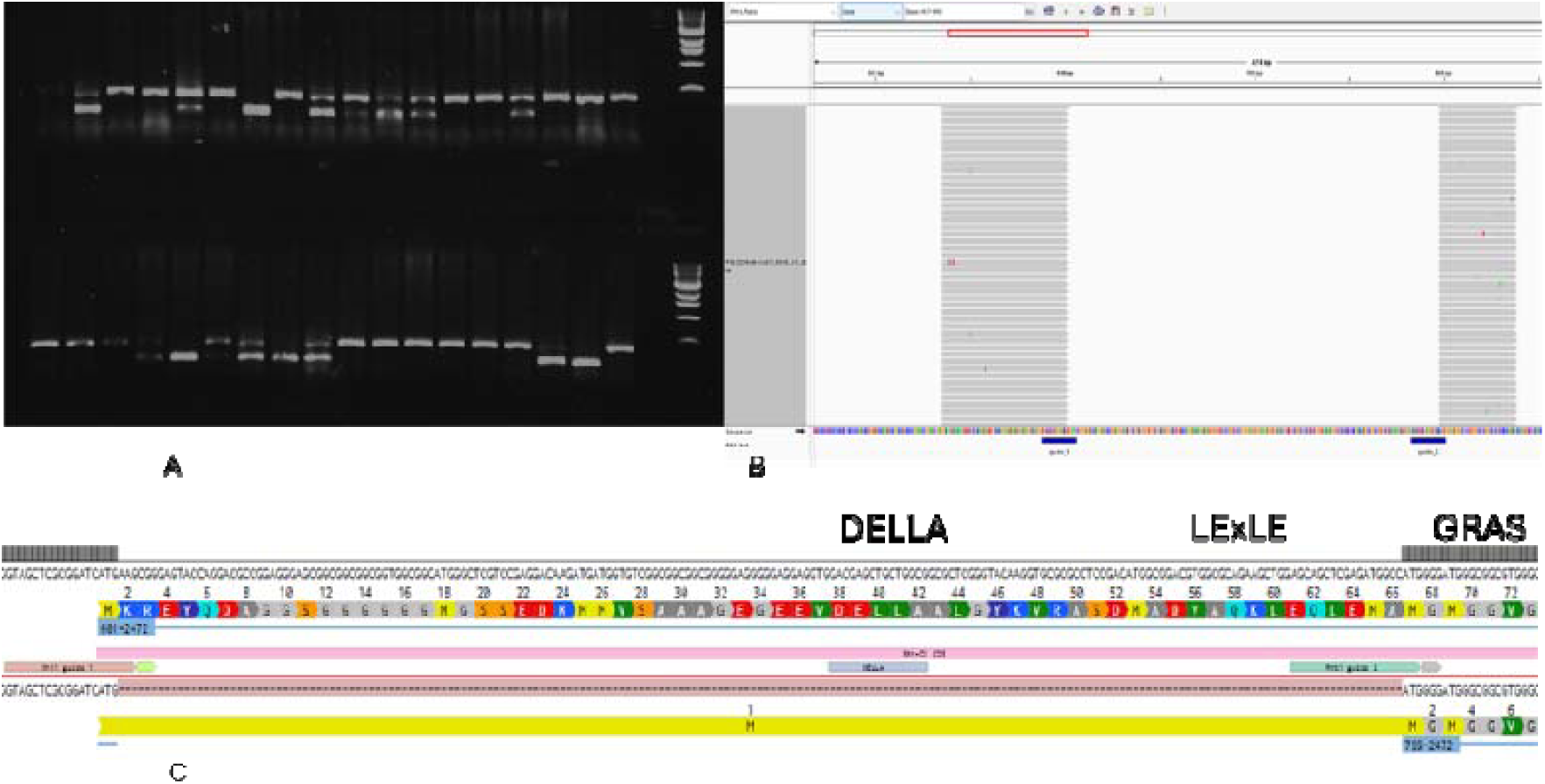
CRISPR/Cas9-mediated deletion of the N-terminal DELLA motif while retaining the C-terminal GRAS domain. (**A**) PCR-based genotyping of representative T₀ plants from the Watkins wheat line WATDE0005 showing putative homozygous (Hom), heterozygous (Het) and wild-type (WT) CRISPR/Cas9 genotypes. Hom plants were identified by the presence of the edited allele only (lower molecular weight band), whereas het plants showed both edited and wild-type alleles (two bands). Non-edited plants retained the wild-type allele only (higher molecular weight band). (**B**) Integrative Genomics Viewer (IGV) visualization of representative CRISPR/Cas9-induced 195-bp deletions identified by targeted amplicon sequencing. (**C**) Schematic representation of the selected homozygous edited allele (113.03.05-CNV2). The in-frame 195-bp deletion removes the DELLA motif while maintaining the downstream C-terminal GRAS domain, generating an N-terminally truncated protein. The edited allele and retained GRAS domain are illustrated using Benchling.

Analysis of targeted amplicon sequencing revealed substantial variation in edited haplotype frequencies among the 166 analysed T₀ plants. Based on the highest-frequency edited haplotype detected in each plant, 80 plants (48.2%) showed high editing frequencies (≥75%), while 36 plants (21.7%) showed intermediate frequencies (50–<75%) and 26 plants (15.7%) showed lower frequencies (30–<50%). A further 10 plants (6.0%) had edited haplotypes below 30%, whereas 14 plants (8.4%) showed no detectable edited haplotype. Overall, 142 of 166 plants (85.5%) contained an edited haplotype at a frequency of ≥30%, with 116 plants (69.9%) showing edited haplotypes at ≥50%.

Haplotype analysis further demonstrated substantial variation in editing outcomes among individual T₀ plants. Among the plants containing detectable editing, 80 plants carried highly enriched predominant edited haplotypes (≥75%), while additional plants showed predominant edited haplotypes at intermediate or lower frequencies. Several plants displayed multiple edited haplotypes within the same individual, indicating considerable allelic diversity generated by CRISPR/Cas9 editing. The broad distribution of edited haplotype frequencies is consistent with the mosaic nature of editing events commonly observed in T₀ plants. Collectively, these results demonstrate high editing activity and substantial allelic heterogeneity at the target locus, with a large proportion of plants carrying highly enriched edited haplotypes suitable for subsequent selection and phenotypic analysis.

At the sequence level, 42 plants with deletions of the DELLA motif while retaining the downstream GRAS domain. Interestingly, transgene copy number analysis identified several plants with no detectable T-DNA (copy number = 0) that nevertheless carried the expected CRISPR/Cas9-induced edits. These plants were therefore classified as transgene-free edited lines, indicating that the editing construct had been eliminated while the edited allele was retained. The recovery of such plants demonstrates the feasibility of generating precisely edited, transgene-free Watkins wheat lines suitable for downstream functional studies and breeding applications.

We identified three primary haplotypes harbouring deletions within the DELLA motif that preserved the downstream, in-frame GRAS domain (Table 4A). In haplotypes 02 and 10, the native start codon was eliminated by single-guide RNA 1 (sgRNA1), leaving the reinitiation start codon to function as the primary start codon. Conversely, haplotype 16 retained the native start codon and featured the intended in-frame tandem methionines (Table 4A). T_0_ individuals exhibiting editing frequencies greater than 75% were selected for subsequent analysis (Table 4B). Plant height varied considerably among T_0_ edited plants within the same Watkins genetic background. Compared with their corresponding non-transformed controls, most edited plants showed reduced height, although others exhibited heights similar to the controls. For example, WATDE0005-derived plants ranged from 32 to 74.9 cm, compared with 90 cm for the control, while WATDE0401-derived plants ranged from 43 to 136 cm, compared with 136 cm for the control (Table 4B). This variation may reflect the chimeric nature of T_0_ plants, in which edited and non-edited cells can coexist within the same plant, potentially resulting in variable phenotypic expression. Among these, WATDE0005 plant ID 113.03.05 was the first T_0_ plant advanced to the T_1_ generation. This line possessed the hap-16, 195-bp deletion (Fig. 4B) that removed the DELLA domain but left the C-terminal GRAS domain intact and in frame (Fig. 4C). Illumina NGS confirmed the deletion, indicating an 88.25% editing frequency in the parental T_0_ plant (Table 4B).

**Table 4.**
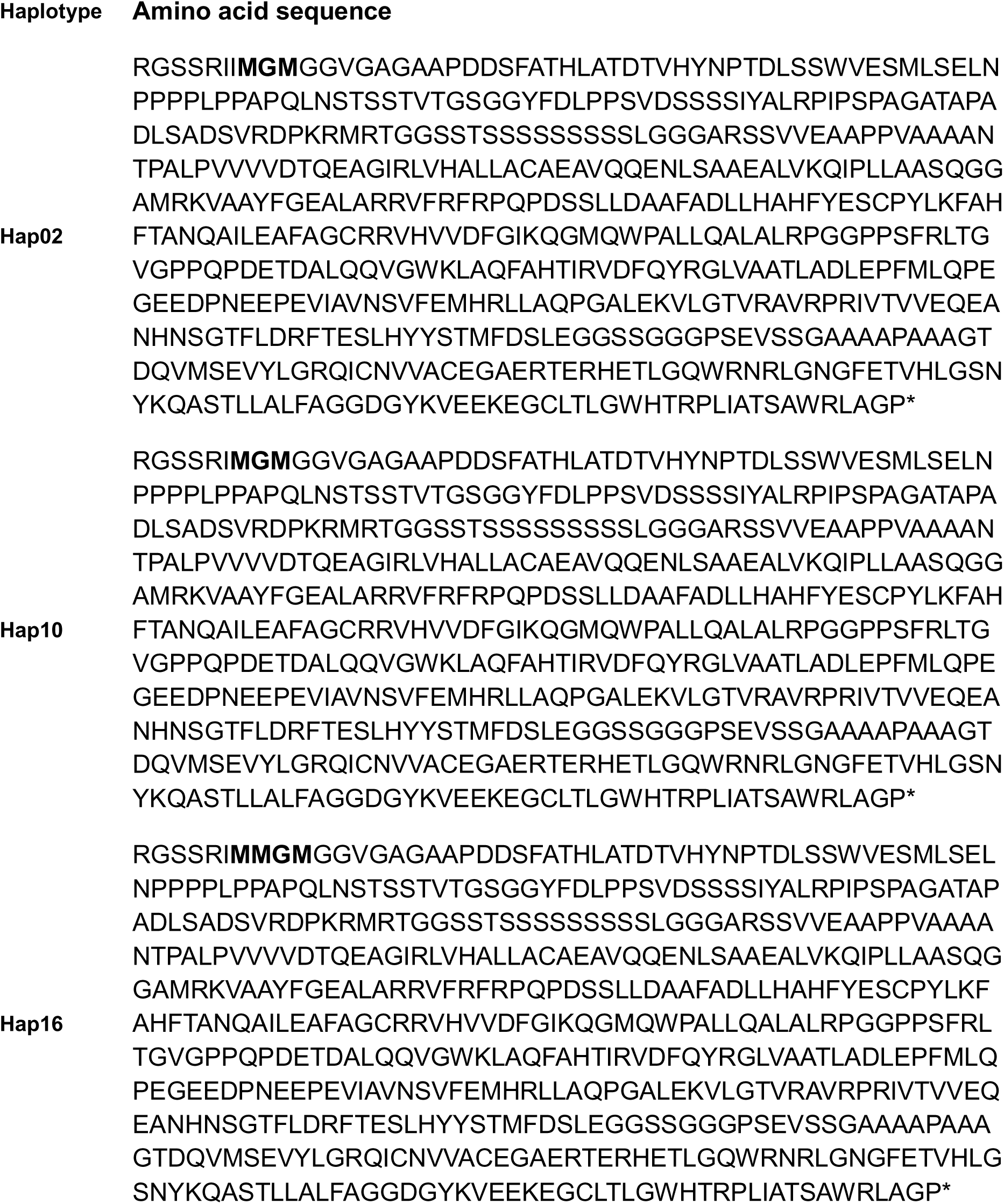
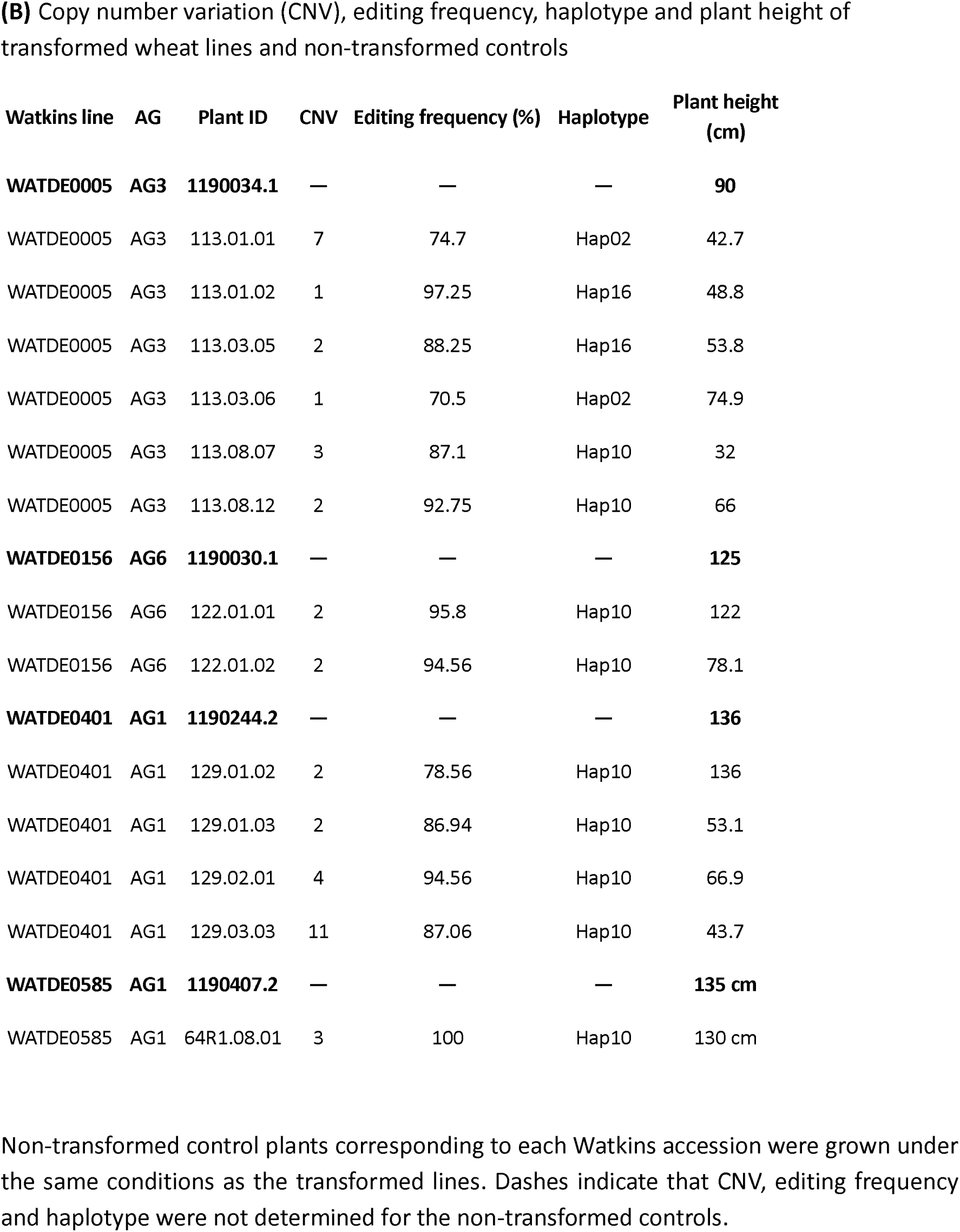
(A) Main in-frame haplotypes carrying deletions of the DELLA motif while retaining the downstream GRAS domain in frame.

The T₀ plant WATDE0005 (line 113.03.05), carrying the in-frame 195-bp deletion (Fig. 5A), was selected for further analysis of the T₁ generation. In figure 5B T₁ plants are presented from left to right as a homozygous edited dwarf plant, a heterozygous edited semi-dwarf plant, and a wild-type (WT) plant with tall phenotype. The progressive reduction in plant height associated with the *Rht-1* genotype demonstrates the phenotypic effect of the targeted DELLA-region modification and further supports the role of the *Rht-1* DELLA region in regulating plant architecture. Copy-number analysis showed that this specific T₀ line contained two copies of the T-DNA. Segregation analysis of the T₁ generation subsequently identified transgene-free edited plants in which the T-DNA had been eliminated while the desired 195-bp deletion was retained. The recovery of these transgene-free edited lines demonstrates the feasibility of generating precisely edited, non-transgenic Watkins wheat germplasm suitable for downstream functional analyses and future breeding applications.

**Figure 5.**
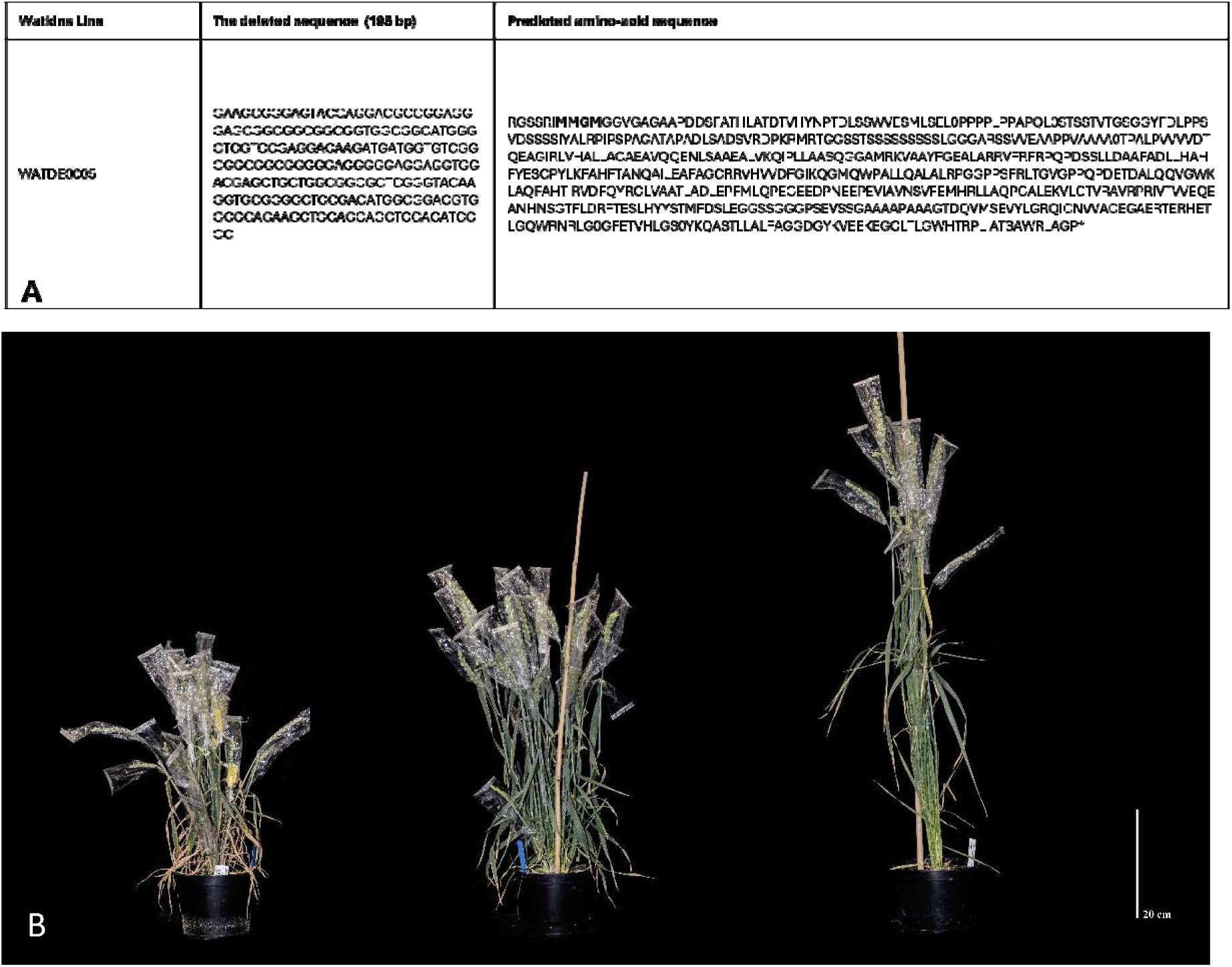
T1 Watkins landrace plants (WATDE0005) with deletion of the N-terminal DELLA domain from the C-terminal GRAS. (**A**) 195-bp deleted nucleotide sequence of the Rht-1 locus. The initiation of translation at the start codon, indicated in bold. (**B**) The edited T_1_ plants were derived from the T_0_ plant 113.03.05 (WATDE0005) showing an 88.25% edited haplotype frequency. Plants are shown from left to right in the following order: homozygous edited plant, displaying a dwarf phenotype; heterozygous edited plant, displaying a semi-dwarf phenotype; and wild-type (WT) plant, displaying a tall phenotype. Zygosity was confirmed by Illumina NGS.

### Cytosine Base Editing with TaCBE6b-intron in Wheat

The major semi-dwarfing alleles *Rht-B1b* and *Rht-D1b* are semi-dominant gain-of-function variants of the homoeologous *Rht-1* genes on chromosomes 4B and 4D, respectively. Each allele confers a similar reduction in plant height, while the presence of both alleles produces an approximately additive effect (Hedden 2003). Given the importance of these alleles in wheat improvement, we sought to introduce a defined C-to-T substitution that generated a premature stop codon on the D-genome *Rht-1* homoeologue. This stop codon was one codon downstream to the naturally occurring stop in *Rht-D1b*, and more representative of the stop codon found in *Rht-B1b*.

To make C to T change to create a premature stop codon on D genome we used a cytosine base editor. The highly active cytosine base editor, CBE6b (Zhang et al., 2024), was employed for targeted base editing in wheat. CBE6b was selected for its enhanced editing efficiency, improved selectivity, and reduced sequence-context dependency compared with earlier TadA-derived cytosine base editors (TadCBEs). The CBE6b editing window spans protospacer positions 3–13 from the 5T end, with maximal activity reported between positions 4 and 8. The target cytosine was positioned within this optimal editing window, enabling efficient introduction of the desired premature stop codon.

To evaluate cytosine base-editing activity in wheat, 25 independent transgenic plants were screened for targeted C-to-T conversion. Four plants showed detectable cytosine base-editing activity, of which two carried the desired nucleotide conversion that introduced a premature stop codon. The edited allele was highly enriched in these plants, with edited haplotype frequencies of 59.3% and 83.0%, respectively (Table 5).

**Table 5.** (A) Amplicon sequencing analysis of individual edited T0 plants showing subgenome-specific base-editing frequencies. (A) Early stop-codon introduction by cytosine base editing. Predicted amino acid (AA) sequences were derived from the nucleotide sequences of plants 106.03.01 and 106.11.02 showing the early stop-codon downstream of the DELLA domain.

| Cultivar | Plant ID | CNV | Mutation | Ref | Edited | A genome (%) | B genome (%) | D genome (%) |
| --- | --- | --- | --- | --- | --- | --- | --- | --- |
| Paragon | 106.03.01 | 4 | SNP | C | T | 0 | 0 | 83 |
| Paragon | 106.11.02 | 4 | SNP | C | T | 0 | 0 | 59.3 |

MKREYQDAGGSGGGGGMGSSSEDKMMVSAAAGEGEEVDELLAALGYKVRASD MADVAQKLE\*

Although earlier TadCBEs enable programmable C•G-to-T•A conversion, they can exhibit residual A•T-to-G•C activity and reduced performance in certain sequence contexts, particularly in GC-rich regions. CBE6b was developed through phage-assisted evolution to overcome these limitations, resulting in improved selectivity and consistent activity across diverse target sequences (Zhang et al. 2024). Originally developed and characterised in mammalian systems, CBE6b was adapted for wheat through codon optimisation and incorporation of seven introns to enhance expression and editing activity. Using the resulting TaCBE6b-intron system, we successfully converted the targeted cytosine residue to thymine, recreating the premature stop codon associated with the *Rht-B1b* Green Revolution semi-dwarf premature-stop variant in the D-genome *Rht-D1* homeologue (Fig. 6A). This additional variant generated using the base-editing approach expands the range of *Rht-1* alleles.

**Figure 6.**
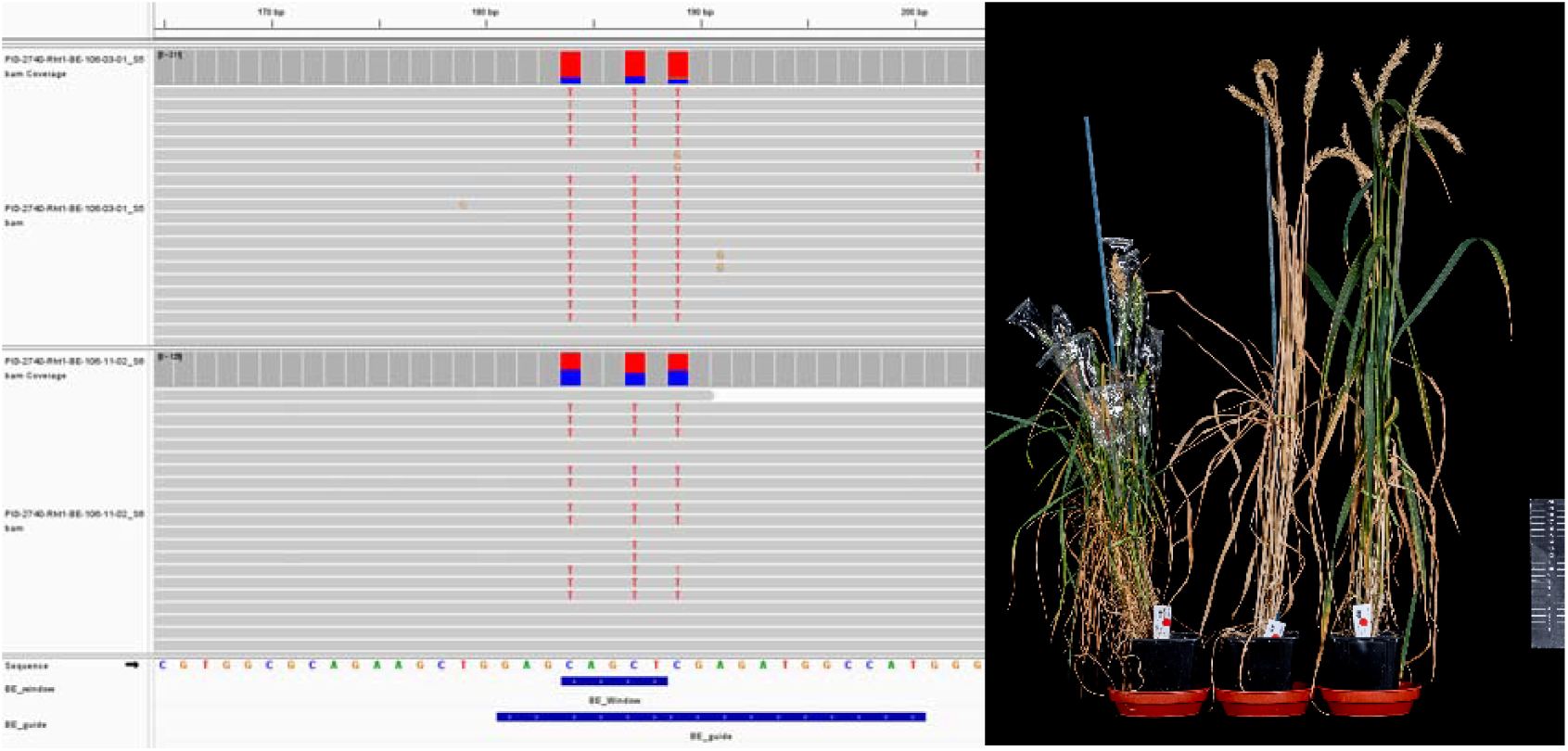
Precise generation of the Rht-B1b Green Revolution semi-dwarf allele through cytosine base editing. **(A)** IGV visualization of targeted amplicon sequencing confirming conversion of the targeted cytosine residue to thymine (C>T substitution), precisely recreating the naturally occurring *Rht-B1b* semi-dwarf allele within the D subgenome in lines 106.03.01 and 106.11.02. **(B)** Phenotypic characterization of edited plants showing the semi-dwarf growth phenotype associated with the engineered *Rht-B1b* allele compared with the non-edited control plants.

In addition to the desired C-to-T conversion generating the premature stop codon, additional cytosine substitutions were detected within the editing window. However, these substitutions occurred downstream of the introduced stop codon and upstream of the reinitiation ATG, these were therefore not predicted to alter the functional outcome of the engineered allele. To further confirm the predicted consequence of the editing event, amino acid (AA) sequences were predicted from the nucleotide sequences of plants 106.03.01 and 106.11.02, in which the edited region was located downstream of the DELLA domain in the D-genome *Rht-D1b* allele. The predicted protein sequences confirmed the introduction of the premature stop codon.

Plants carrying the engineered allele displayed the expected semi-dwarf growth phenotype, confirming the functional impact of the base-edited mutation (Fig. 6B). Notably, the premature stop codon generated in the present study is located one position downstream of the naturally occurring mutation and therefore represents an additional *Rht-D1* allelic variant.

### Prime Editing Enables Targeted Introduction of the *Rht-D1b* Allele in Watkins Wheat Landraces

The precise introduction of the *Rht-D1b* allele requires a G-to-T nucleotide substitution that is not accessible using conventional cytosine or adenine base-editing chemistries. Prime editing provides a broader editing scope by enabling precise installation of defined nucleotide substitutions and therefore provides an alternative approach for engineering *Rht-1* alleles that cannot be generated using base editors. Following the successful implementation of gene editing approaches in Watkins wheat landraces, we evaluated the enhanced plant prime editing system, ePPEplus (Ni et al., 2023), for precise modification of *Rht-D1*. Based on its broader editing flexibility and the higher editing efficiencies observed in our initial experiments, prime editing was subsequently prioritised for the targeted engineering of *Rht-1* alleles in Watkins wheat landraces.

The ePPEplus construct was designed to introduce the semi-dwarf *Rht-D1b* allele through precise conversion of the endogenous wild-type nucleotide to the desired edited sequence. The *Rht-D1b* allele contains a premature UAG stop codon within the *Rht-D1* coding sequence, resulting in the production of a truncated DELLA protein with reduced sensitivity to GA-mediated degradation. The premature stop codon is located one codon upstream of the corresponding stop codon in *Rht-B1b*. The first prime-editing experiment targeting *Rht-D1* demonstrated successful allele conversion across multiple Watkins genetic backgrounds, establishing ePPEplus as an effective platform for precise nucleotide engineering in diverse wheat germplasm.

Targeted amplicon sequencing of 53 primary transgenic (T₀) plants identified 17 plants carrying G-to-T conversion events above 30% of the NGS reads (Table 6A). Edited haplotypes represented 30.8 to 100% of sequencing reads among positive individuals (Table 6B). The highest edited haplotype frequencies were observed in plants derived from WATDE0394, with an individual plant 80R2.03.02 reaching 100% edited haplotype frequency in the D-genome *Rht-D1* amplicon (Fig. 7A). This plant line was advanced to the T_1_ generation, where 100% of the progeny (92/92) were homozygous for the edit according to KASP marker analysis (Table 6C). Several additional plants also showed high levels of prime editing, including edited haplotype frequencies of 98.2%, 96.4%, 51.8% and 47.0%, demonstrating substantial prime-editing activity in individual T₀ plants.

**Figure 7.**
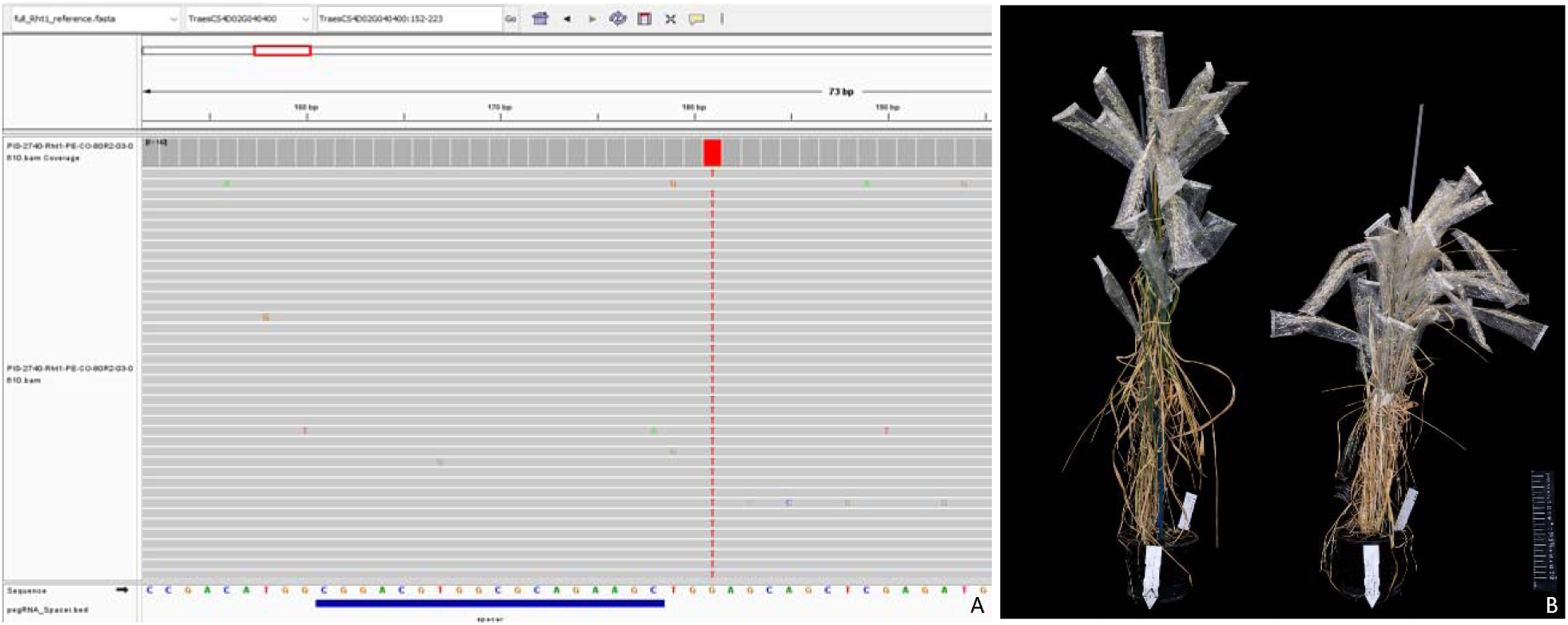
**Generation of the semi-dwarf *Rht-D1b* allele through prime editing in T_₀_ wheat.** (**A**) IGV visualization of targeted amplicon sequencing from T₀ plant WATDE0394 (80R2.03.02), confirming prime editing-mediated installation of the designed G-to-T nucleotide substitution at the Rht-D1 locus. (**B**) Phenotypic characterization of the prime-edited WATDE0394 plant (80R2.03.02) Wildtype to the left, prime edited to the right showing semi dwarf phenotype.

**Table 6.** Prime editing efficiency and haplotype frequencies of edited *Rht-D1* alleles in Watkins wheat landraces.

| Watkins line | Ancestral Group (AG) | No. of plants analysed | Percentage of edited plants (%) |
| --- | --- | --- | --- |
| WATDE0394 | AG3 | 7 | 100.0 |
| WATDE0549 | AG4 | 2 | 100.0 |
| WATDE0639 | AG7 | 3 | 100.0 |
| WATDE0005 | AG3 | 5 | 0.0 |
| WATDE0585 | AG1 | 3 | 0.0 |
| WATDE0385 | AG3 | 9 | 0.0 |
| WATDE0008 | AG7 | 7 | 0.0 |
| WATDE0052 | AG7 | 1 | 0.0 |
| WATDE0706 | AG1 | 10 | 50.0 |
| WATDE0588 | AG3 | 4 | 0.0 |
| WATDE0051 | AG7 | 2 | 0.0 |

| Watkins line | Plant ID | CNV | Mut. | Ref | Ed. | A (%) | B (%) | D (%) | Height (cm) |
| --- | --- | --- | --- | --- | --- | --- | --- | --- | --- |
| WATDE0394 | 1190240-2 | — | — | — | — | — | — | — | 131 |
| WATDE0394 | 80R2.03.02 | 2 | SNP | G | T | 0 | 0 | 100 | 69 |
| WATDE0394 | 80R2.03.03 | 2 | SNP | G | T | 0 | 0 | 45.62 | 70 |
| WATDE0394 | 80R2.03.04 | 2 | SNP | G | T | 0 | 0 | 51.84 | 85 |
| WATDE0394 | 80R2.03.06 | 2 | SNP | G | T | 0 | 0 | 96.4 | 74 |
| WATDE0394 | 80R2.03.07 | 2 | SNP | G | T | 0 | 0 | 98.2 | 71 |
| WATDE0394 | 80R2.03.08 | 3 | SNP | G | T | 0 | 0 | 33.7 | 75 |
| WATDE0394 | 80R2.03.11 | 2 | SNP | G | T | 0 | 0 | 39.12 | 72 |
| WATDE0549 | 1190375-1 | — | — | — | — | — | — | — | 140 |
| WATDE0549 | 81R1.02.01 | 70 | SNP | G | T | 0 | 39.2 | 47 | 51–85 |
| WATDE0549 | 81R1.02.04 | 10 | SNP | G | T | 42.16 | 0 | 44.12 | 78 |
| **WATDE0639** | **1190453-1** | — | — | — | — | — | — | — | **138** |
| WATDE0639 | 83R2.01.01 | 2 | SNP | G | T | 0 | 0 | 34.53 | 97 |
| WATDE0639 | 83R2.02.01 | 2 | SNP | G | T | 0 | 0 | 30.81 | 67–93 |
| WATDE0639 | 83R2.02.02 | 9 | SNP | G | T | 0 | 0 | 34.47 | 71–103 |
| **WATDE0706** | **1190516-1** | — | — | — | — | — | — | — | **102** |
| WATDE0706 | 196.03.01 | 1 | SNP | G | T | 52.1 | 61.8 | 50.7 | 47–85 |
| WATDE0706 | 196.04.01 | 1 | SNP | G | T | 48.6 | 0 | 59.47 | 53–65 |
| WATDE0706 | 196.04.02 | 1 | SNP | G | T | 36.34 | 0 | 39.06 | 46–67 |
| WATDE0706 | 196.06.03 | 3 | SNP | G | T | 0 | 63.88 | 10.84 | Died |
| WATDE0706 | 196.09.04 | 1 | SNP | G | T | 26.98 | 43.25 | 58.84 | 35–63 |

| Population | G/G | G/T | T/T | Total |
| --- | --- | --- | --- | --- |
| 80R2.03.02 | 92 | 0 | 0 | 92 |
| 81R1.02.04 | 14 | 56 | 26 | 96 |
| 83R2.02.01 | 0 | 14 | 1 | 15 |
| 196.09.04 | 26 | 49 | 21 | 96 |

Prime-editing activity was also detected, >30% NGS reads, in additional Watkins landraces, including WATDE0549, WATDE0639 and WATDE0706, demonstrating that ePPEplus activity was not restricted to a single genetic background. Editing frequencies varied substantially among genotypes, ranging from undetectable levels to complete conversion. Edited individuals were recovered from lines WATDE0394, WATDE0549, WATDE0639, and WATDE0706, where 100%, 100%, 100%, and 50% of analysed transformed plants showed prime editing, respectively, whereas no edited individuals were recovered from several other transformed genotypes (Table 6A). This variation highlights the influence of genetic background on prime-editing performance in wheat and is consistent with the genotype dependence observed for other gene editing technologies.

In our DELLA deletion and base editing experiments, we successfully targeted the D genome by exploiting SNP differences unique to the *Rht-1D* homeolog; conversely, the prime editing pegRNA protospacer had to be designed in a conserved region upstream of the required G-to-T prime edit and therefore targeted all three *Rht-1* homeologs. Notably, despite the simultaneous targeting of all three homeologs, the *Rht-1D* homeolog was edited with noticeably higher efficiency than its A and B counterparts (Table 6B). This discrepancy in prime editing efficiency among the *Rht-1* homeologs could be attributed to variations in local chromatin structure and subgenome-specific epigenetic landscapes (Kim et al. 2024). It is highly probable that the *Rht-1D* locus resides within a more open, euchromatic region, thereby facilitating enhanced pegRNA binding and increased accessibility of the prime editing complex compared to the more tightly wound or heterochromatic environments of the *Rht-1A* and *Rht-1B* loci (Kim et al. 2024; Ramírez-González et al. 2018). The premature stop codons introduced via prime editing into *Rht-1A* and *Rht-1B* represent novel, unique variants. Notably, the induced stop codon within *Rht-1B* is located exactly one codon upstream of the mutation responsible for the naturally occurring, semi-dwarfing *Rht-B1b* allele. Furthermore, because this represents the first report of a premature stop codon introduced into the *Rht-1* A-genome homeolog, we propose designating this novel allele as *Rht-A1b*.

The enhanced editing activity observed with ePPEplus is consistent with the system improvements reported by Ni et al. (2023). The ePPEplus incorporates an engineered reverse transcriptase containing the R221K, V223A and N394K substitutions, additional nuclear localisation signals and a wheat codon-optimised coding sequence to enhance nuclear accumulation and editing activity. In addition, incorporation of the Csy4 RNA-processing system facilitates pegRNA maturation and stability, contributing to improved prime-editing efficiency (Liu et al., 2021; Ni et al., 2023).

Importantly, the engineered *Rht-D1b* allele was associated with the expected semi-dwarf phenotype in the T₀ plants analysed (Table 6B). A proportion of the T_0_ plants exhibiting distinct height strata and the values represented in table 6B represent the maximum height of each stratum. This phenotypic variability likely reflects T_0_ spatial chimerism, in which a mosaic of prime-edited and unedited cells within individual plant sectors results in differing height strata. All prime-edited plants showed a reduction in height compared to non-transformed control. In WATDE0394 line 80R2.03.02, the edited plant reached approximately 69 cm in height compared with 131 cm for the corresponding non-transformed control (Fig. 7B). Similar reductions in plant height were observed in other edited Watkins backgrounds, including WATDE0549 and WATDE0639, where edited plants reached mean heights of 71.3 (SD=18.0) and 86.2 (SD= 16.2) cm, respectively, compared with 140 and 138 cm for their corresponding non-transformed controls. The most pronounced effect was observed in WATDE0706, where T_0_ edited plants had a mean height 57.6 (SD= 15.6) cm compared with 102 cm for the non-transformed control. This pronounced reduction in height seen in WATDE0706 may reflect additional contributions from A and B subgenome editing (Table 6B). Together, these results indicate that installation of the engineered *Rht-D1b* nucleotide substitution was associated with reduced plant height across multiple genetic backgrounds.

T₁ progeny were further genotyped using KASP markers to confirm inheritance of the engineered *Rht-D1b* allele, and PCR analysis was performed to identify transgene-free edited plants. Four lines were selected for further T₁ analysis. Line 80R2.03.02 was homozygous for the engineered allele, whereas lines 81R1.02.04, 83R2.02.01, and 196.09.04 were heterozygous in the T₀ generation (Table 6B). A total of 299 T₁ plants from these four prime-editing populations were screened for segregation of the engineered allele. Among these plants, 132 were homozygous positive (G/G), 119 were heterozygous (G/T), and 48 were negative (T/T) wild type (Table 6C). Homozygous, heterozygous, and negative plants were selected for further phenotypic analysis, including GA₃ treatment, to assess the effects of the engineered *Rht-D1b* allele (Fig. 8). A two-way (Type III) ANOVA revealed a significant genotype × treatment interaction (*F*_2,42_ = 5.13, 0.0101) indicating that the response to gibberellic acid (GA) treatment depended on the plant genotype. Consequently, we evaluated the treatment effects within each group using Tukey’s HSD post-hoc test. Under mock conditions, growth did not differ significantly among the three genotypes (*p* = 0.790; all sharing the letter “a” in Fig. 8C). Conversely, GA treatment induced highly significant differences in growth across genotypes (*p* < 0.0001). Under GA exposure, wild-type (WT) plants exhibited the greatest mean growth (25.0 cm), followed by heterozygous (19.0 cm) and homozygous mutants (13.6 cm) (Fig. 8C).

**Figure 8.**
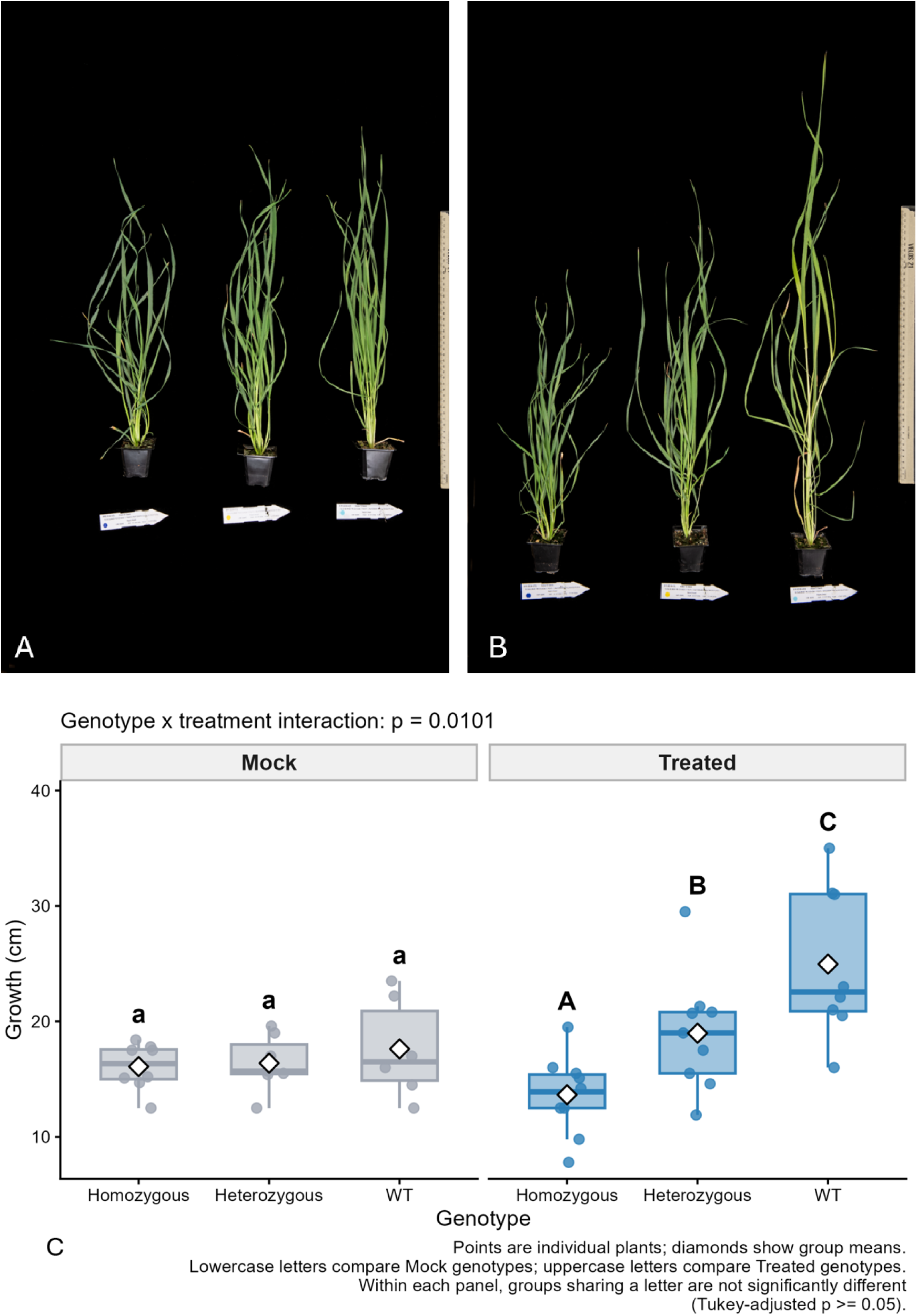
GA responsiveness of prime-edited *Rht-D1b* plants. Representative prime-edited plants are shown from left to right as homozygous (hom), heterozygous (het), and wild-type (WT) genotypes. **(A)** Mock-treated controls were sprayed with water containing a small amount of ethanol. **(B)** GA -treated plants show a clear dosage-dependent reduction in GA -induced growth, with homozygous plants exhibiting the strongest reduction and heterozygous plants displaying an intermediate response. This pattern is consistent with the semi-dominant effect of the reduced-height *Rht-1* phenotype. **(C)** Under Mock conditions, growth did not differ significantly among the three genotypes ( 0.790); hence, all Mock groups share the letter “a”. Under GA treatment, genotype had a highly significant effect ( 0.0001). WT had the greatest mean growth (25.0 cm), followed by Heterozygous plants (19.0 cm) and Homozygous plants (13.6 cm). All three treated genotypes differed significantly from each other according to Tukey’s test, represented by the letters A, B and C.

These results demonstrate a clear dosage-dependent reduction in GA₃ responsiveness associated with the engineered *Rht-D1b* allele. Homozygous plants showed the strongest reduction in GA₃-induced growth, while heterozygous plants displayed an intermediate response, consistent with the semi-dominant nature of the *Rht-1* reduced-height phenotype.

Collectively, these results demonstrate that prime editing can introduce defined nucleotide substitutions into *Rht-D1* in diverse wheat landrace backgrounds. The ability to install nucleotide changes that are inaccessible to conventional base editors highlights prime editing as a complementary technology for precision engineering of agronomically important alleles in genetically diverse wheat germplasm.

## Conclusion

This study demonstrates that advanced genome-editing technologies can be successfully deployed across genetically diverse Watkins wheat landraces. By combining an efficient *Agrobacterium*-mediated transformation system with developmental regulator-assisted regeneration, we demonstrate robust CRISPR/Cas9 mutagenesis, cytosine base editing and prime editing across diverse wheat genetic backgrounds.

CRISPR/Cas9-mediated editing achieved high efficiencies across Watkins accessions, while the TaCBE6b cytosine base editor was, to our knowledge, applied for the first time in plants and optimised for wheat, enabling precise creation of a novel D-genome *Rht-D1* allele. The engineered *Rht-D1* allele carries a premature stop codon positioned one nucleotide downstream of the stop codon in the naturally occurring *Rht-D1b* allele. We have also created new novel Rht1 alleles on the A and B subgenomes using prime editing. Importantly, prime editing enabled precise installation of the Rht-D1b allele, with edited haplotypes reaching frequencies of up to 100% in positive T_0_ plants, demonstrating the potential of optimized prime editing for precise allele engineering in wheat. Together, these approaches extend advanced genome-editing technologies beyond the model spring wheat cultivar ‘Fielder’ and provide complementary strategies for introducing defined nucleotide changes, engineered alleles and novel functional variants without relying exclusively on conventional double-strand break-mediated mutagenesis in genetically diverse wheat landraces.

Targeted engineering of the Rht-1 loci generated a diverse allelic series, including frameshift mutations, precise DELLA motif deletions and engineered semi-dwarfing alleles. In particular, deletion of the conserved DELLA motif while maintaining the downstream GRAS domain in-frame provides an experimental framework for dissecting the contribution of the N-terminal regulatory region to DELLA stability, gibberellin responsiveness and growth repression. These engineered alleles therefore provide valuable resources for investigating the molecular basis of Rht-1-mediated plant architecture across diverse genetic backgrounds.

The Watkins Collection provides an exceptional reservoir of genetic diversity, including alleles associated with disease resistance, abiotic stress tolerance and other agronomically important traits. Introducing defined semi-dwarfing alleles directly into these backgrounds enables the combination of Green Revolution growth traits with previously underutilised ancestral diversity, while retaining the broader genetic architecture and potentially favourable native alleles of the landraces.

Collectively, the transformation and genome-editing platforms established here provide a robust framework for precision engineering of diverse wheat germplasm. The combination of efficient transformation with CRISPR/Cas9, cytosine base editing and increasingly efficient prime editing enables both functional dissection of gene function and targeted generation of breeding-relevant alleles. These resources provide a foundation for expanding precision breeding into genetically diverse wheat germplasm and for developing novel founder lines for future breeding and composite cross populations.

## Acknowledgments

The authors acknowledge and thank Phil Robinson of JIC Photography Department for photographs and Mark Youles of TSL SynBio for supplying some of the L0 golden gate components. This research formed part of the core project of the Wheat Genetic Improvement Network (WGIN) which is supported by a commission by the Department for the Environment, Food and Rural Affairs (Defra, C24770).

## References

1. Anzalone AV, Randolph PB, Davis JR, Sousa AA, Koblan LW, Levy JM, Chen PJ, Wilson C, Newby GA, Raguram A, Liu DR (2019) Search-and-replace genome editing without double-strand breaks or donor DNA. Nature 576 (7785):149–157. doi:10.1038/s41586-019-1711-4

2. Bilgic H, Hakki EE, Pandey A, Khan MK, Akkaya MS (2016) Ancient DNA from 8400 Year-Old Çatalhöyük Wheat: Implications for the Origin of Neolithic Agriculture. PLOS ONE 11 (3):e0151974. doi:10.1371/journal.pone.0151974

3. Borlaug NE (1968) Wheat breeding and its impact on world food supply. Australian Academy of Science Australia,

4. Chen PJ, Liu DR (2023) Prime editing for precise and highly versatile genome manipulation. Nat Rev Genet 24 (3):161–177. doi:10.1038/s41576-022-00541-1

5. Cheng S, Feng C, Wingen LU, Cheng H, Riche AB, Jiang M, Leverington-Waite M, Huang Z, Collier S, Orford S, Wang X, Awal R, Barker G, O’Hara T, Lister C, Siluveru A, Quiroz-Chávez J, Ramírez-González RH, Bryant R, Berry S, Bansal U, Bariana HS, Bennett MJ, Bicego B, Bilham L, Brown JKM, Burridge A, Burt C, Buurman M, Castle M, Chartrain L, Chen B, Denbel W, Elkot AF, Fenwick P, Feuerhelm D, Foulkes J, Gaju O, Gauley A, Gaurav K, Hafeez AN, Han R, Horler R, Hou J, Iqbal MS, Kerton M, Kondic-Spica A, Kowalski A, Lage J, Li X, Liu H, Liu S, Lovegrove A, Ma L, Mumford C, Parmar S, Philp C, Playford D, Przewieslik-Allen AM, Sarfraz Z, Schafer D, Shewry PR, Shi Y, Slafer GA, Song B, Song B, Steele D, Steuernagel B, Tailby P, Tyrrell S, Waheed A, Wamalwa MN, Wang X, Wei Y, Winfield M, Wu S, Wu Y, Wulff BBH, Xian W, Xu Y, Xu Y, Yuan Q, Zhang X, Edwards KJ, Dixon L, Nicholson P, Chayut N, Hawkesford MJ, Uauy C, Sanders D, Huang S, Griffiths S (2024) Harnessing landrace diversity empowers wheat breeding. Nature 632 (8026):823–831. doi:10.1038/s41586-024-07682-9

6. Debernardi JM, Tricoli DM, Ercoli MF, Hayta S, Ronald P, Palatnik JF, Dubcovsky J (2020) A GRF-GIF chimeric protein improves the regeneration efficiency of transgenic plants. Nat Biotechnol 38 (11):1274–1279. doi:10.1038/s41587-020-0703-0

7. Develtere W, Waegneer E, Debray K, De Saeger J, Van Glabeke S, Maere S, Ruttink T, Jacobs TB (2023) SMAP design: a multiplex PCR amplicon and gRNA design tool to screen for natural and CRISPR-induced genetic variation. Nucleic Acids Research 51 (7):e37–e37. doi:10.1093/nar/gkad036

8. Evans LT (1998) Feeding the ten billion. Plants and population growth john r walton 128

9. Grützner R, Martin P, Horn C, Mortensen S, Cram EJ, Lee-Parsons CWT, Stuttmann J, Marillonnet S (2021) High-efficiency genome editing in plants mediated by a Cas9 gene containing multiple introns. Plant Commun 2 (2):100135. doi:10.1016/j.xplc.2020.100135

10. Han L, Wang X, Benke R, Tibbs-Cortes LE, Zhao P, Sanguinet KA, Zhang Z, Xu S, Yu J, Li X (2025) Integrated phenomic and genomic analyses unveil modes of altered phenotypic plasticity during wheat improvement. Genome Biol 26 (1):256. doi:10.1186/s13059-025-03740-1

11. Hayta S, Smedley MA, Bayraktar M, Forner M, Backhaus A, Lister C, Clarke M, Uauy C, Griffiths S (2026) Controlling GRF4-GIF1 expression for efficient, genotype-independent transformation across wheat cultivars. The Plant Journal 125 (6):e70799. 10.1111/tpj.70799

12. Hedden P (2003) The genes of the Green Revolution. Trends in Genetics 19 (1):5–9. doi:10.1016/S0168-9525(02)00009-4

13. Jin S, Lin Q, Gao Q, Gao C (2023) Optimized prime editing in monocot plants using PlantPegDesigner and engineered plant prime editors (ePPEs). Nat Protoc 18 (3):831–853. doi:10.1038/s41596-022-00773-9

14. Kim S, Yuan JB, Woods WS, Newton DA, Perez-Pinera P, Song JS (2024) Corrigendum: Chromatin structure and context-dependent sequence features control prime editing efficiency. Frontiers in Genetics Volume 15–2024. doi:10.3389/fgene.2024.1391923

15. Lazo GR, Stein PA, Ludwig RA (1991) A DNA transformation-competent Arabidopsis genomic library in Agrobacterium. Bio/technology (Nature Publishing Company) 9 (10):963–967

16. Liu Y, Yang G, Huang S, Li X, Wang X, Li G, Chi T, Chen Y, Huang X, Wang X (2021) Enhancing prime editing by Csy4-mediated processing of pegRNA. Cell Res 31 (10):1134–1136. doi:10.1038/s41422-021-00520-x

17. Ramírez-González RH, Borrill P, Lang D, Harrington SA, Brinton J, Venturini L, Davey M, Jacobs J, van Ex F, Pasha A, Khedikar Y, Robinson SJ, Cory AT, Florio T, Concia L, Juery C, Schoonbeek H, Steuernagel B, Xiang D, Ridout CJ, Chalhoub B, Mayer KFX, Benhamed M, Latrasse D, Bendahmane A, Consortium IWGS, Wulff BBH, Appels R, Tiwari V, Datla R, Choulet F, Pozniak CJ, Provart NJ, Sharpe AG, Paux E, Spannagl M, Bräutigam A, Uauy C, Korol A, Sharpe AG, Juhász A, Rohde A, Bellec A, Distelfeld A, Akpinar BA, Keller B, Darrier B, Gill B, Chalhoub B, Steuernagel B, Feuillet C, Chaudhary C, Uauy C, Pozniak C, Ormanbekova D, Xiang D, Latrasse D, Swarbreck D, Barabaschi D, Raats D, Sergeeva E, Salina E, Paux E, Cattonaro F, Choulet F, Kobayashi F, Keeble-Gagnere G, Kaur G, Muehlbauer G, Kettleborough G, Yu G, Šimková H, Gundlach H, Berges H, Rimbert H, Budak H, Handa H, Small I, Bartoš J, Rogers J, Doležel J, Keilwagen J, Poland J, Melonek J, Jacobs J, Wright J, Jones JDG, Gutierrez-Gonzalez J, Eversole K, Nilsen K, Mayer KFX, Kanyuka K, Singh K, Gao L, Concia L, Venturini L, Cattivelli L, Spannagl M, Mascher M, Hayden M, Abrouk M, Alaux M, Luo M, Valárik M, Benhamed M, Singh NK, Sharma N, Guilhot N, Ravin N, Stein N, Olsen O-A, Gupta OP, Khurana P, Chhuneja P, Bayer PE, Borrill P, Leroy P, Rigault P, Sourdille P, Hernandez P, Flores R, Ramirez-Gonzalez RH, King R, Knox R, Appels R, Zhou R, Walkowiak S, Galvez S, Biyiklioglu S, Nasuda S, Sandve S, Chalabi S, Weining S, Sehgal S, Jindal S, Belova T, Letellier T, Wicker T, Tanaka T, Fahima T, Barbe V, Tiwari V, Kumar V, Tan Y (2018) The transcriptional landscape of polyploid wheat. Science 361 (6403):eaar6089. doi:doi:10.1126/science.aar6089

18. Ross JJ (1994) Recent advances in the study of gibberellin mutants. Plant Growth Regulation 15 (3):193–206. doi:10.1007/BF00029892

19. Scalzitti N, Kress A, Orhand R, Weber T, Moulinier L, Jeannin-Girardon A, Collet P, Poch O, Thompson JD (2021) Spliceator: multi-species splice site prediction using convolutional neural networks. BMC Bioinformatics 22 (1):561. doi:10.1186/s12859-021-04471-3

20. Smedley MA, Hayta S, Clarke M, Harwood WA (2021) CRISPR-Cas9 Based Genome Editing in Wheat. Current Protocols 1 (3):e65. 10.1002/cpz1.65

21. Würschum T, Langer SM, Longin CF (2015) Genetic control of plant height in European winter wheat cultivars. Theor Appl Genet 128 (5):865–874. doi:10.1007/s00122-015-2476-2

22. Youssefian S, Kirby E, Gale M (1992) Pleiotropic effects of the GA-insensitive Rht dwarfing genes in wheat. 2. Effects on leaf, stem, ear and floret growth. Field Crops Research 28 (3):191–210

23. Zhang E, Neugebauer ME, Krasnow NA, Liu DR (2024) Phage-assisted evolution of highly active cytosine base editors with enhanced selectivity and minimal sequence context preference. Nature Communications 15 (1):1697. doi:10.1038/s41467-024-45969-7

