## Supplementary Figure 1 for "Integrating Green Revolution *Rht-1* Alleles in Diverse Wheat Landraces Using CRISPR/Cas9, Cytosine Base Editing and Prime Editing"

**GGTCTCAAATGGTTCACTGCCGTATAGGCAGGGCGGACGTGGCGCAGAAGCGTTTAAGAGCTATGCTGGAAACAGCATAGCAAGTTTAAATAAGGCTAGTCCGTTATCAACTTGAAAAAGTGGCACCGAGTCGGTGCTCGAGCTGCTACAGCTTCTGCGCCACGCGGTTCTATCTAGTTACGCGTTAAACCAACTAGAAGTTCACTGCCGTATAGGCAGGGATCCGCTTTGAGACC**

**Csy4 processing site (red)**

**Rht guide RNA (light blue)**

**Guide Scaffold (purple)**

**Reverse transcription template (orange)**

**Prime Binding Site (green)**

**RNA Knot tevopreQ1 (dark yellow)**

**Supplementary Figure 1.** Nucleotide sequence of the pegRNA used for prime editing of *Rht-1*. The Csy4 processing site, *Rht-D1* guide RNA, guide RNA scaffold, reverse-transcription template (RTT), prime-binding site (PBS), and RNA knot tevopreQ1 are indicated by red, light blue, purple, orange, green, and dark yellow, respectively.
